# HIV-1 preintegration complex preferentially integrates the viral DNA into nucleosomes containing trimethylated histone 3-lysine 36 modification

**DOI:** 10.1101/2022.02.16.480507

**Authors:** Nicklas Sapp, Nathaniel Burge, Khan Cox, Santosh Thapa, Prem Prakash, Muthukumar Balasubramaniam, Devin Christenson, Min Li, Jared Linderberger, Mamuka Kvaratskhelia, Jui Pandhare, Robert Craigie, Wesley Sundquist, Michael G Poirier, Chandravanu Dash

**Author notes:** Corresponding Author Chandravanu Dash, PhD Old Hospital Bldg-CAHDR, Room 5027, 1005 Dr. DB Todd Jr Blvd., Nashville, TN 37208, USA.

## Abstract

HIV-1 DNA integration into the host chromosomes is carried out by the preintegration complex (PIC). The PIC contains the viral DNA, virally encoded integrase enzyme and other critical viral/host factors. The PIC-associated viral DNA is preferentially integrated into gene bodies of actively transcribing genes. Here, we identify a biochemical mechanism underlying the preference of PIC-mediated viral DNA integration (PIC-VDI). Specifically, we observed that the PIC-VDI into human chromatin is preferred over the genomic DNA. Surprisingly, nucleosome core particles without any histone modifications were not preferred for PIC-VDI when compared to the analogous naked DNA. However, PIC-VDI was markedly enhanced with nucleosomes containing the trimethylated histone 3 lysine 36 (H3K36me3), an epigenetic mark linked to HIV-1 DNA integration preference. Interestingly, we observed that nucleosomes with flanking linker DNA promoted PIC-VDI in the presence of LEDGF/p75. We also discovered that nucleosomes with linker DNA and H3K36me3 served as the optimal substrate for PIC-VDI. Mapping of the integration sites within these substrates identified preference of specific regions of the nucleosome core DNA for integration. Finally, we provide biochemical and genetic evidence that histone H1 protein, that condenses the chromatin, negatively regulates HIV-1 DNA integration, consistent with the integration preference for *open chromatin* structure. Collectively, these results identify the role of specific chromatin marks that drive HIV-1 integration preference and define the optimal substrate requirement for efficient DNA integration by the PIC.

## Introduction

HIV-1, the causative agent of AIDS, has infected approximately 80 million people worldwide, with ∼38 million people living with the virus currently (<u>UNAIDS, 2021</u>). Potent antiretroviral therapy (ART) has rendered HIV-1 infection a chronic disease and have significantly reduced the disease burden to AIDS (<u>UNAIDS, 2021</u>). However, ART is not curative and life-long ART can cause severe toxicity and comorbid conditions ^1^. Therefore, a clear understanding of the mechanism of HIV-1 replication is key for developing novel and improved therapeutic strategies. HIV-1 infection is dependent on the integration of the reverse transcribed viral DNA into host chromosomes ^2–4^. This critical step is carried out by the virally encoded integrase (IN) enzyme in a multistep process ^5^. First, IN carries out the 3’-processing of the viral DNA ends and then via a strand-transfer step, the viral DNA is inserted into host genomic DNA ^6–9^. This intricate process of DNA integration is carried out by the viral preintegration complex (PIC) which is composed of IN, the viral DNA, and other viral and host factors ^10–15^. Accordingly, establishment of the proviral genome is required for generation of progeny virions and ultimately establishment of viral reservoirs, which remains the principal barrier to HIV-1 cure ^16–18^.

HIV-1 integration is a non-random event that targets specific genic regions of human chromosomes. There is strong evidence that HIV-1 DNA is integrated into hot spots in the genome. While protein-coding genes account for less than 2% of the genome, they represent over half of the detected HIV-1 integration sites ^19–25^. Analysis of HIV-1 integration sites reveals consistent genic integration patterns in *open chromatin* across both cell lines and patient derived samples ^23, 24, 26, 27^. These preferred regions for integration are characterized by the active transcription of highly expressed genes, gene density, and epigenetic factors characteristic of *open chromatin* ^21, 25–31^ . The chromosomal context of proviral integration is important, as these genomic characteristics influence proviral transcription ^32, 33^. To carry out viral DNA integration into the genomic hot spots, the HIV-1 PIC must engage and overcome the structural and functional barriers of chromosomal landscape. A single copy of the human genome can extend over two meters, yet it is packaged into a nucleus with an average diameter of 10 *µm* ^34^. This enormous task of genome compaction is achieved by organizing the DNA into a nucleoprotein polymer called chromatin. The basic repeating unit of chromatin is a nucleosome, each of which contains a nucleosome core composed of an octamer of histone proteins (H2A, H2B, H3 and H4) that wraps 1.6 turns of a 147 bp of DNA ^34, 35^. The nucleosome core is connected to the adjacent nucleosome core by a segment of linker DNA. The linker DNA is often associated with the histone protein (H1) that further compacts the genome ^36–38^. In addition to these genomic organization mechanisms, an array of epigenetic modifications on the histone proteins and DNA, add further complexity to the structural and functional landscape of the chromatin ^39–41^. Therefore, specific mechanisms within the nuclear environment are required for the PIC to successfully insert the viral DNA into the hot spots.

Despite the targeting of HIV-1 DNA into genic regions for integration, the contribution of chromatin to HIV-1 integration has been elusive. Nevertheless, there is evidence that retroviral INs, including that of HIV-1, specifically target the DNA wrapped within nucleosomes for integration ^42–46^. Notably, retroviral INs preferentially target the outward facing major groove of the nucleosomal DNA ^44–47^. The distortion of DNA as it bends around the histone octamer is thought to facilitate integration into the nucleosomes. Interestingly, retroviral INs differ in how they integrate viral DNA within chromatinized DNA templates ^48–54^. While HIV and MLV target nucleosome cores, some reports show that nucleosome arrays are less preferred for integration when compared to analogous naked DNA^50, 53, 55^. In contrast, ALV IN activity is specifically enhanced by nucleosome compaction^48^. Evidently, host proteins such as LEDGF/p75, BET, and others also play key roles in viral DNA integration, especially with chromatin substrates ^48–50, 53, 55– 58^. Particularly, the interaction between HIV-1 IN and LEDGF/p75 is linked to integration site-selection to genes and can specifically stimulate HIV-1 IN strand-transfer reactions *in vitro* ^59–70^ .

Still, the molecular and biochemical details underlying retroviral DNA integration in the context of chromatin are not fully understood. Here we report a biochemical mechanism for the preference of the HIV-1 DNA integration into *open chromatin*. For this we used HIV-1 PICs extracted from infected T-cell lines and examined its ability to integrate into isolated chromatin. Analysis of PIC-mediated integration with biochemically assembled nucleosomes without histone tail modifications showed reduced integration. Notably, the addition of a tri-methylated histone tail resulted in enhanced PIC-associated viral DNA integration into nucleosomes. Addition of linker DNA to nucleosomes modulated the effect of LEDGF/p75 on HIV-1 integration. Furthermore, chromatin compaction by the linker histone H1^0^ negatively regulated HIV-1 integration. Finally, through sequencing analysis, integration preference was identified within specific regions of the nucleosomal DNA. Overall, our study provides critical biochemical evidence for HIV-1 PIC-mediated integration preference into *open chromatin*.

## Results

### Chromatin is the preferred substrate for HIV-1 PIC-mediated viral DNA integration

HIV-1 infection is dependent on integration of the viral DNA into host chromosomes by the pre-integration complex (PIC). It is well-established that HIV-1 DNA is preferentially integrated into the gene body of dense, actively transcribing genes ^19, 20, 24, 30^. However, the biochemical determinants of integration site preference within the chromatin landscape are not fully understood. To better understand the mechanism of HIV-1 integration into chromatin, we adopted a method that measured PIC-associated viral DNA integration (PIC-VDI) using 300 ng of target substrates from isolated chromatin DNA, de-chromatinized genomic DNA, biochemically reconstituted nucleosome core particles (NCP) with or without linker DNA and the analogous DNA sequences. HIV-1 PICs extracted from acutely infected SupT1 cells were competent for viral DNA integration *in vitro* ^71–74^ (SI-Fig. 1A-B). The PIC-VDI assay is specific since the HIV-1 integrase inhibitor raltegravir (RAL) significantly inhibited the DNA integration activity (SI-Fig. 1B). Chromatin, isolated from HEK293T cells, contained the canonical histone proteins, and lacked detectable levels of cytoplasmic protein markers (Fig. 1A-B and SI-Fig. 1C). Importantly, micrococcal nuclease digestion of the isolated chromatin resulted in a “ladder-like” pattern with the smallest DNA band at 150 base pairs (bp) (Fig. 1C and SI-Fig. 13C), equivalent to the length of DNA wrapped around a nucleosome ^34, 35, 75^, suggesting optimal protection of the DNA in our chromatin preparation. In parallel, de-chromatinized DNA obtained by deproteination of the chromatin was used as the analogous genomic DNA (gDNA) substrate (Fig. 1C). To probe the preference of PIC-VDI into these substrates, PICs were incubated with either the chromatin or gDNA preparations containing equivalent amount of DNA and PIC-VDI was measured by *alu*- based nested qPCR ^72–74^. Results from these measurements revealed that PIC-VDI levels were significantly higher with the chromatin substrate compared to the isolated gDNA (Fig. 1D-E). Since the chromatin substrate contains the host factors and histone modifications that are positively associated with HIV-1 integration (SI-Fig. 1C), the preference of PIC-VDI into the chromatin is likely driven by host factors that may influence HIV-1 integration and the structural elements of within the nucleosomes.

**Figure 1.**
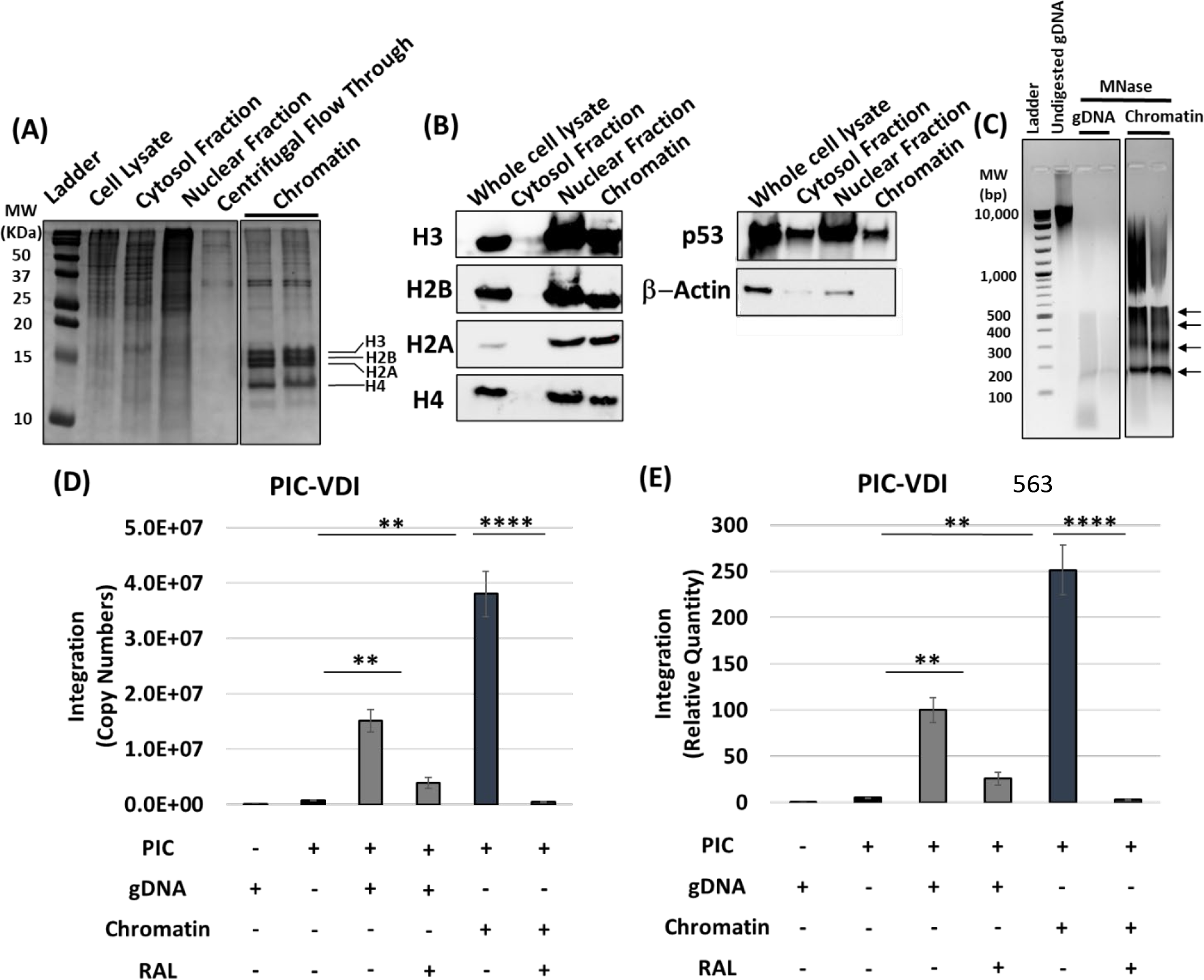
Chromatin is the preferred substrate for HIV-1 PIC-VDI. Chromatin was isolated from HEK293T cells and assessed for the histone proteins and DNA. (A) Fractions from various steps of the chromatin preparation were analyzed by a 15% acrylamide gel, visualized by Coomassie staining. The chromatin fraction containing the canonical histone proteins: H3, H2B, H2A, and H4 are shown. (B) The histone proteins were detected by western blot in the whole cell lysate, cytosol fraction, nuclear fraction, and chromatin (left panel). Additionally, cytoplasmic, and nuclear proteins were probed in each fraction (right panel). (C) Protection of the chromatin DNA within nucleosomes was assessed by micrococcal nuclease (MNase) digestion. The arrows to the right of the gel indicative discrete DNA bands of nucleosome mediated protection. (D) HIV-1 PIC-VDI was measured by nested-PCR and represented as the copy numbers of integrated viral DNA, using 300 ng of chromatin and deproteinated genomic DNA as targets. As a negative control, 1 µM RAL-the integrase strand transfer inhibitor was used. (E) The copies of HIV-1 DNA integration were plotted relative to the integration into genomic DNA. The results in E are shown as the mean of viral DNA copy numbers of at least three replicates, with the error bars indicated the standard error mean (SEM). * Represents P > 0.05, ** P = 0.01 to 0.05, *** P = 0.01 to 0.001, P = 0.001 to 0.0001, and **** P < 0.0001.

### Nucleosome core particles (NCP) without histone tail modifications are not preferred for PIC-VDI

There is evidence that nucleosomes are the preferred substrates of retroviral integration ^43–46, 49, 76^. Therefore, to better understand the enhanced PIC activity with chromatin, we probed PIC-VDI using biochemically assembled NCPs ^77^. NCPs were assembled using the Widom 601 nucleosome positioning sequence (NPS) DNA ^78^ and human histone octamers assembled by a stepwise approach using purified recombinant histone H2A, H2B, H3, and H4 (Fig. 2A). The 147 bp NPS-DNA (Fig. 2B) was added to the assembled histone octamer (Fig. 2C) and formation of the NCP was confirmed by an electromobility shift assay (EMSA) (Fig. 2D). Then, 300 ng of the nucleosome core particle and the nucleosome positioning sequence were used as substrates to measure PIC-VDI by nested qPCR using specific primers that amplify the junctions of integrated viral DNA into the target substrate. Surprisingly, PIC-VDI into the NPS- DNA was significantly higher when compared to the NCP (Fig. 2E). Specificity of these assays were demonstrated by the dramatically reduced PIC-VDI by 1 µM RAL in the presence of both the NCP and NPS (Fig. 2E). There are a number of key differences that could explain why these results deviate from the prior studies showing that nucleosomes are preferred substrates for integration compared to either naked DNA within the same chromatinized plasmid or analogous DNA. One difference is that our study of nucleosome substrates for retroviral DNA integration utilizes HIV-1 PICs compared to the *in vitro* studies of purified HIV-1, MLV, and PFV integrase enzyme ^43–46, 48–50, 55, 58^. The PIC associated factors present in our PICs may influence integration preference beyond what might be driven by integrase alone. Further, the published studies used nucleosomes, chromatinized substrates and histones derived from cellular sources ^34, 48–50, 53, 55, 79^, whereas our studies use strongly positioned, biochemically assembled recombinant nucleosome.

**Figure 2.**
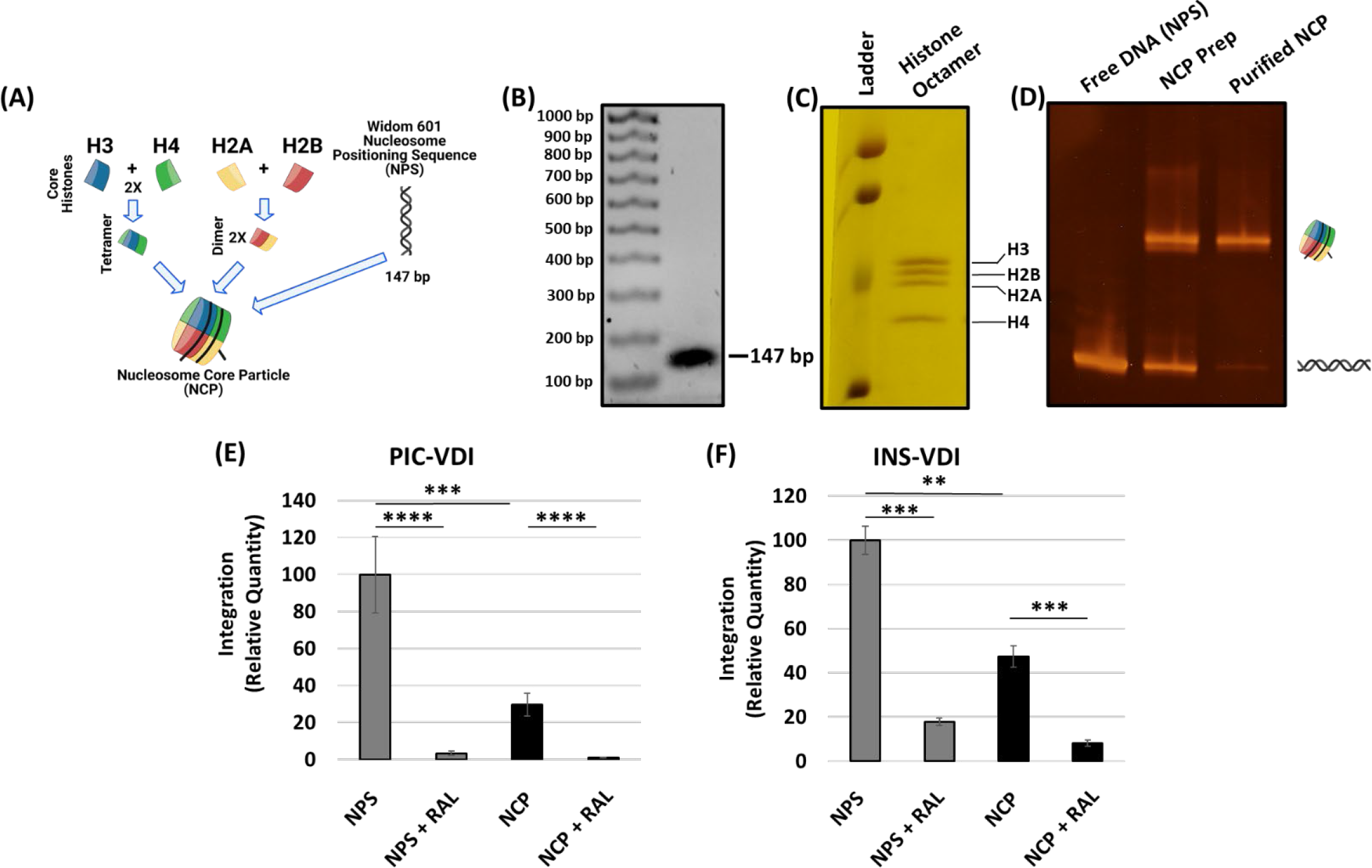
The NCP is a barrier to HIV-1 VDI. (A) A schematic of the biochemical assembly of nucleosome core particle (NCP). The NCP is assembled with the Widom 601 nucleosome positioning sequence (NPS) and a recombinant human histone octamer. The histone octamer was assembled first by an equimolar addition of histone protein dimers H2A and H2B with the H3/H4 tetramer. (B) The 147 bp NPS DNA used for nucleosome assembly was analyzed on an 1.5% agarose gel and visualized by ethidium bromide staining. (C) The presence of individual histone proteins in the octameric histone assembly were visualized by SDS-AGE and Coomassie staining. (D) Biochemical assembly of the nucleosome was analyzed by an electro-mobility shift assay (EMSA). (E) PIC-VDI was assessed using 300 ng of the NPS and the NCP as targets, with 1 µM of RAL serving as a specific inhibitor for HIV-1 integration. (F) The intasome (INS) VDI was measured using quantitative PCR with the NPS and the NCP serving as targets. The data is represented as the relative quantity of VDI in reference to the NPS and error bars were generated from the SEM of at least three independent experiments. * Represents P > 0.05, ** P = 0.01 to 0.05, *** P = 0.01 to 0.001, P = 0.001 to 0.0001, and **** P < 0.0001.

Prior studies have indicated that the Widom nucleosome position sequence used in our study forms more stable nucleosomes than native DNA sequences^76, 78, 80^.

Since we observed reduced PIC-VDI with the NCP substrate, we probed whether this lack of substrate preference is PIC specific. To test this, we used HIV-1 intasomes (INS), which are biochemically assembled complexes of purified integrase with viral DNA sequences from the long terminal repeat (LTR) region ^81, 82^. We used the purified recombinant IN containing a Ssod7 domain for solubility ^82^ and viral DNA mimics (25 bp and 27 bp) (SI-Fig. 2B). We tested INS-mediated viral DNA integration (INS-VDI) by a qPCR-based assay to amplify the integration junctions into these specific target substrates (SI-Fig. 2B). This qPCR-based approach selectively amplified the viral and target DNA junction only when both the INS and target substrate were added to the reaction mixture (SI-Fig. 2C). Since RAL dramatically reduced the INS-VDI, the assay specifically measures DNA integration (SI-Fig. 2C). Finally, INS-VDI was compared with the NPS and NCP substrates in these assay conditions (Fig. 2F). Surprisingly, VDI was significantly higher with the NPS substrate when compared to the NCP (Fig. 2F), an observation similar to the PIC-VDI (Fig. 2E). Interestingly, the level of reduction in INS-VDI into NCP substrates was ∼50% compared to a marked reduction of ∼70% in PIC-VDI, suggesting that NPCs confer an additional barrier for PICs to insert the viral DNA. Taken together these observations suggest that NCPs are not preferred for HIV-1 DNA integration by the PIC or the INS.

### Trimethylated NCPs at Histone 3 Lysine 36 enhance PIC-VDI

In the chromatin, nucleosomes are decorated with chemical modifications primarily in the N-terminal tails of the histones ^40, 83, 84^. Given our unexpected results that NCPs without histone modifications are not preferred for PIC-VDI, we assessed whether NCPs containing specific histone tail modifications are required for optimal PIC-VDI. One of the histone modifications commonly associated with HIV-1 integration preference is the trimethylated histone 3 at lysine 36 (H3K36me3) ^21, 25, 26, 29– 32, 51, 85^. Therefore, we tested whether the H3K36me3 mark in the nucleosome plays a direct role in HIV-1 integration preference. To test this, we biochemically assembled NCPs with the biochemical mimetic (H3K36Cme3) of the H3K36me3 modification generated by an alkylation reaction (Fig. 3A). The H3K36Cme3 NCP does not enhance nucleosomal DNA unwrapping ^86^ and is biochemically indistinguishable from the post-translational modification (PTM) found in cells ^87^. Our western blot analysis also confirmed the detection of H3K36me3 in isolated chromatin and H3K36Cme3 modified NCPs (Fig. 3B). When we tested PIC-VDI using the H3K36Cme3-NCP as the substrate, we observed a robust increase in PIC-VDI when compared to the analogous unmodified NCP substrate (Fig. 3C-D). Interestingly, PIC-VDI with the H3K36Cme3-NCP substrate was significantly less than the PIC-VDI with the NPS substrate (Fig. 3D). To further probe this paradox, we measured INS-VDI using the NPS, NCP and H3K36Cme3-NCP substrates. In stark contrast to the PIC-VDI (Fig. 3C), INS-VDI with the H3K36Cme3-NCP substrate showed no measurable difference relative to the unmodified NCP (Fig. 3E). However, the INS-VDI was significantly higher with the NPS substrate when compared to both of the NCP substrates (Fig. 3F). These observations suggest that the biochemical determinants of HIV-1 integration in the context of PIC are specific in recognizing the histone H3K36me3 modification in the nucleosome compared to the INS.

**Figure 3.**
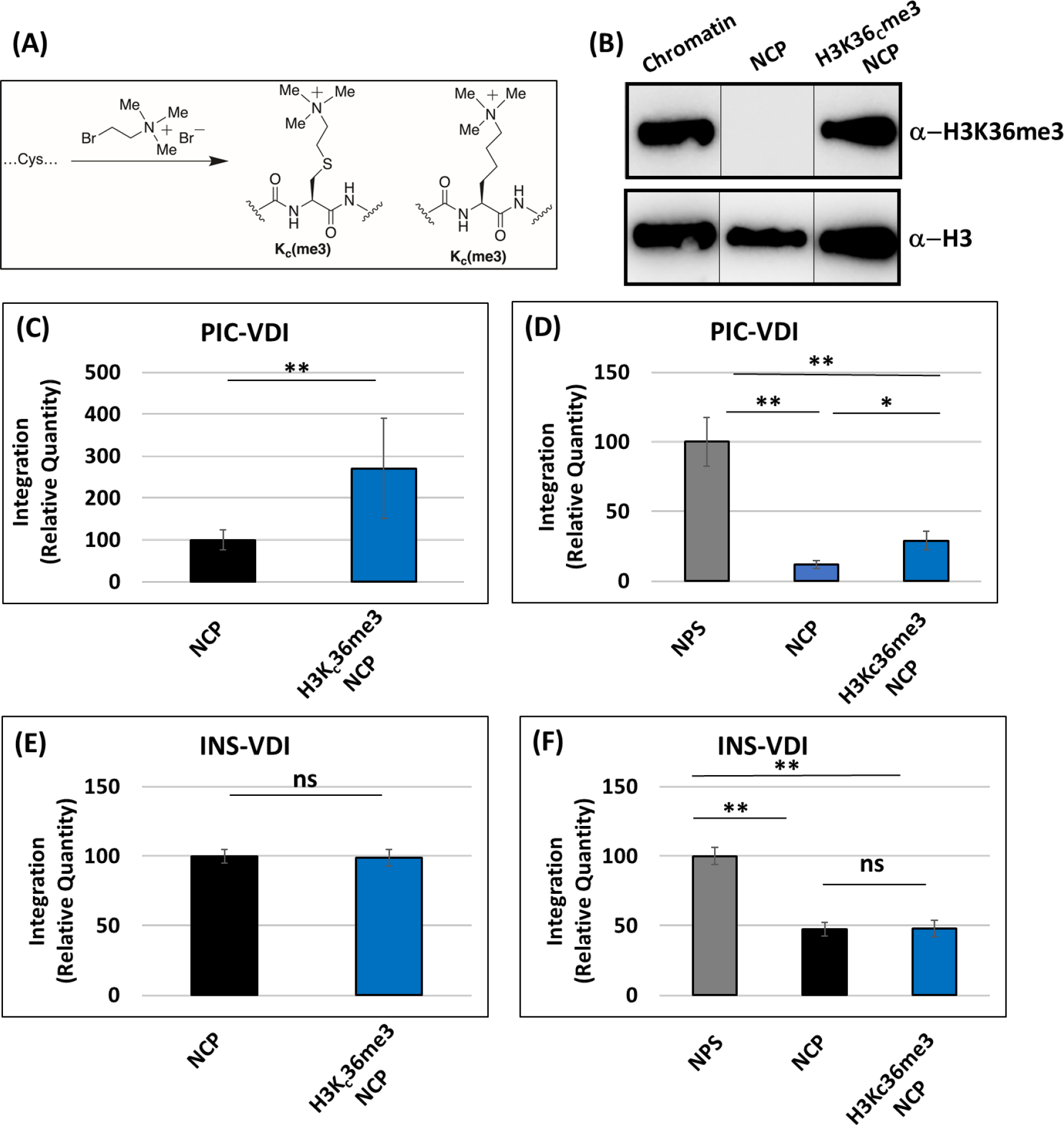
Nucleosomes containing trimethylated NCPs at Histone 3 Lysine 36 enhance PIC- VDI. (A) A schematic illustrating the insertion of the H3K36me3 by site-directed mutagenesis of the K36 to cysteine (K36C). The K36C H3 undergoes alkylation to functionalize the K36C to create a biochemical mimic of the histone 3 trimethylation (H3KC36me3). (B) Western blot analysis of the H3K36me3 and histone H3 in the H3K36Cme3-NCP compared to the unmodified NCP and the isolated chromatin. (C) PIC-VDI was measured with the NCP and the H3K36Cme3-NCP substrates. The integration data was plotted relative to the PIC-VDI into the NCP. (D) The PIC- VDI with the NPS, NCP and H3K36me3-NCP and the data was analyzed in reference to the NPS. (E) INS-VDI was measured comparing the NCP and H3K36me3-NCP. (F) Then the INS-VDI was assessed comparing the NPS, NCP and H3K36me3-NCP. The error bars were determined by the SEM of at least three independent replicates. * Represents P > 0.05, ** P = 0.01 to 0.05, *** P = 0.01 to 0.001, P = 0.001 to 0.0001, and **** P < 0.0001

### LEDGF/p75 addition stimulates INS-VDI but reduces PIC-VDI

In addition to specific histone modifications, the PIC utilizes several host proteins to efficiently integrate the viral DNA into the host chromosomes ^57, 88–91^. Early studies identified LEDGF/p75 as the key host protein to support HIV-1 integration ^59–62, 67–69, 79^. LEDGF/p75 binds to the integrase catalytic core domain (CCD) and stimulates the strand-transfer activity of lentiviral integrases, including HIV, with naked and chromatinized substrates ^92–94^. Therefore, we tested the effects of LEDGF/p75 on INS-VDI and PIC-VDI using nucleosome and naked DNA substrates. First, increasing amounts of purified LEDGF/p75 was supplemented to the INS-VDI reaction mixture containing the NPS and NCP substrates. We observed a dose dependent enhancement of INS-VDI with the NPS-DNA with increasing concentrations of LEDGF/p75 (Fig. 4A). Similarly, LEDGF/p75 addition also stimulated INS-VDI with the NCP (Fig. 4B). Surprisingly, increasing amounts of LEDGF/p75 did not enhance PIC-VDI with the NPS substrate albeit a slight but non-significant decrease at the highest concentrations (Fig. 4C). However, LEDGF/p75 addition significantly reduced PIC-VDI with the NCP (Figure 4D) in contrast to the enhanced effect of the protein in INS-VDI assays (Fig. 4A-B). Notably, our western blot analysis showed that endogenous LEDGF/p75 is present in our PIC preparation (Fig. 4E). Therefore, it is plausible that the PIC-associated endogenous LEDGF/p75 levels is sufficient for VDI and additional amounts of the protein has an inhibitory effect. To test this, we used a fluorescence resonance energy transfer (FRET) based binding assay that utilizes a 5’-Cy3 labeled NPS reconstituted into an NCP containing a Cy5 label on the octamer ^95, 96^. The creative positioning of these two fluorophores allowed for the measurement of NCP accessibility of a test protein near the DNA entry/exit location. Particularly, since LEDGF/p75 is known to bind the NCP at the entry/exit site ^86, 97, 98^, this assay is uniquely suited for our experiment. In this assay, a *GAL4* binding sequence was cloned into the NPS (at the nucleotide positions of 6 to 25) to determine GAL4 accessibility to the NCP in direct competition with LEDGF/p75 (SI-Fig. 4A). Initially, we demonstrated that LEDGF/p75 binds to the NCP by EMSA (SI-Fig. 4B) consistent with the published studies ^97, 98^. Moreover, in the absence of GAL4, addition of increasing amounts of LEDGF/p75 did not alter the FRET efficiency (SI-Fig. 4C). These results established that LEDGF/p75 addition minimally disrupts the interactions between the DNA and histones within the NCP. From our fluorescence polarization analysis, we observed that LEDGF/p75 binds to both the NCP and H3K36Cme3 NCP substrates with a comparable affinity of 30 and 28 nM, respectively (Figure 4F). Then we tested the NCP accessibility to GAL4 in the presence of LEDGF/p75 (Fig. 4G-H). Interestingly, LEDGF/p75 addition reduced the accessibility of GAL4 to the NCP by ∼2-fold. Similarly, GAL4 accessibility to the DNA in the NCP containing H3K36me3 was also reduced by LEDGF/p75. Collectively, these results suggest that LEDGF/p75 can physically or sterically block trans-factors from gaining access to the DNA within the NCP. These data would suggest that excess LEDGF/p75 reduces PIC-VDI by blocking access of PIC-associated viral DNA to the NCP-bound target DNA.

**Figure 4.**
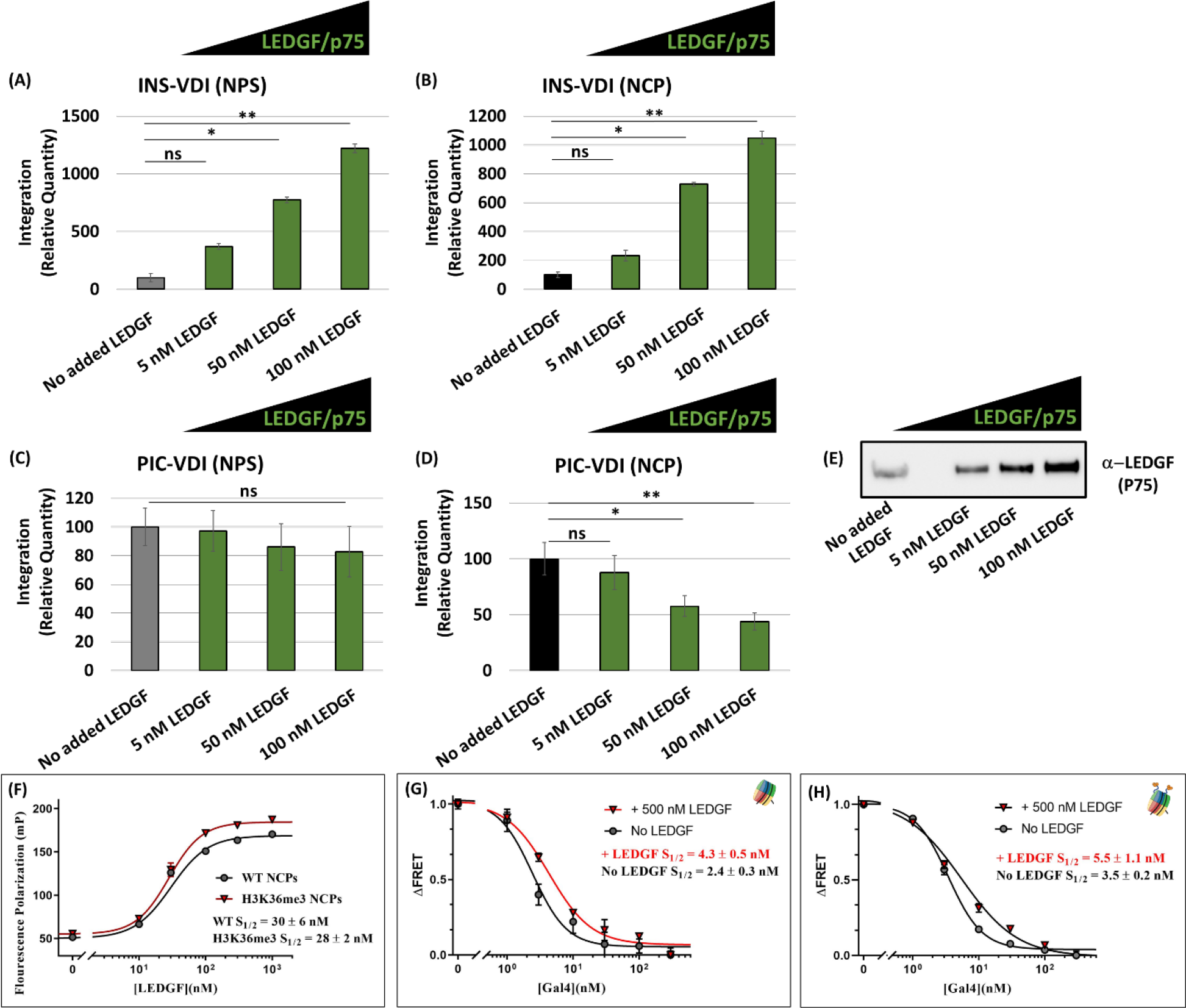
Presence of LEDGF/p75 stimulates INS-VDI but reduces PIC-VDI. (A-B) INS-VDI [25 nM] was measured in the presence of the integration cofactor LEDGF/p75. LEDGF was added to the indicated reaction mixtures at 5, 10 and 25 nM to both the NPS substrate and the NCP substrate. (C-D) PIC-VDI was measured in the presence of LEDGF/p75 with both the NPS substrate and the NCP substrate. (E) Western blot analysis for LEDGF/p75 in the isolated PIC preparation and also in PICs supplemented with the recombinant LEDGF/p75 protein. (F) Fluorescence polarization assay were performed with fluoresceine labeled NCP to determine the LEDGF binding kinetics (S1/2) to the nucleosomes. (G-H) The ensemble FRET measurements were detected by Cy3-Cy5 NCPs (Cy3 on the end of the NPS and Cy5 at the H2AK119C) and with titrations of GAL4 in the presence of LEDGF/P75 at saturating amounts of the unmodified NCP and H3K36Cme3-NCP. The error bars represent the SEM of at least three independent experiments. * Represents P > 0.05, ** P = 0.01 to 0.05, *** P = 0.01 to 0.001, P = 0.001 to 0.0001, and **** P < 0.0001.

### NCPs containing H3K36me3 and linker DNA promote PIC-VDI

Our results show that the NCP is less preferred for both PIC-VDI and INS-VDI compared to the analogous NPS-DNA (Fig. 2E-F). To better understand this surprising result, we tested the effect of the NCP on viral DNA integration by adding a 50 bp linker DNA flanking both ends of the NCP-bound DNA (Fig. 5A-B, SI-Fig 5A), which is more representative of nucleosomes in chromatin ^75, 99^. First, we measured INS-VDI with the 247 bp linker NPS-DNA and compared it with the linker NCP (Fig. 5C). We observed a reduction in the integration with the linker NCP compared to the linker NPS, similar to our results with NPS and NCP substrates (Fig. 2). Furthermore, INS-VDI with the linker- NCP and the linker H3K36Cme3-NCP substrates showed that the histone trimethylation has a negligible effect on viral DNA integration (Fig. 5D). Results from INS-VDI with all of three linker substrates again showed the preference of INS-VDI for the NPS substrate (Fig. 5E), demonstrating that naked DNA is a highly preferred substrate for the HIV-1 INS. Finally, we tested PIC-VDI with the linker NPS-DNA and NCP. The linker NPS was preferred for PIC-VDI over the linker NCP (Fig. 5F), similar to the data of INS-VDI (Fig. 5C-E) and PIC-VDI with non-linker DNA containing substrates (Fig. 3C-D). However, the results of PIC-VDI with the linker NCP and linker H3K36Cme3-NCP revealed a significant enhancement in PIC-VDI with the H3K36Cme3-NCP substrate (Fig. 5G). Most importantly, comparative analysis of PIC-VDI with all of the linker substrates showed that the linker H3K36Cme3-NCP supported significantly higher VDI than the linker NCP and linker NPS-DNA (Fig. 5H). This is in contrast to the results with INS-VDI (Fig. 5E), suggesting that the combination of the H3K36me3 and the presence of linker DNA provide a biochemically favored environment specifically for HIV-1 PIC-VDI. These results provide the first biochemical evidence for the H3K36me3-mediated stimulation of HIV-1 integration by the PIC.

**Figure 5.**
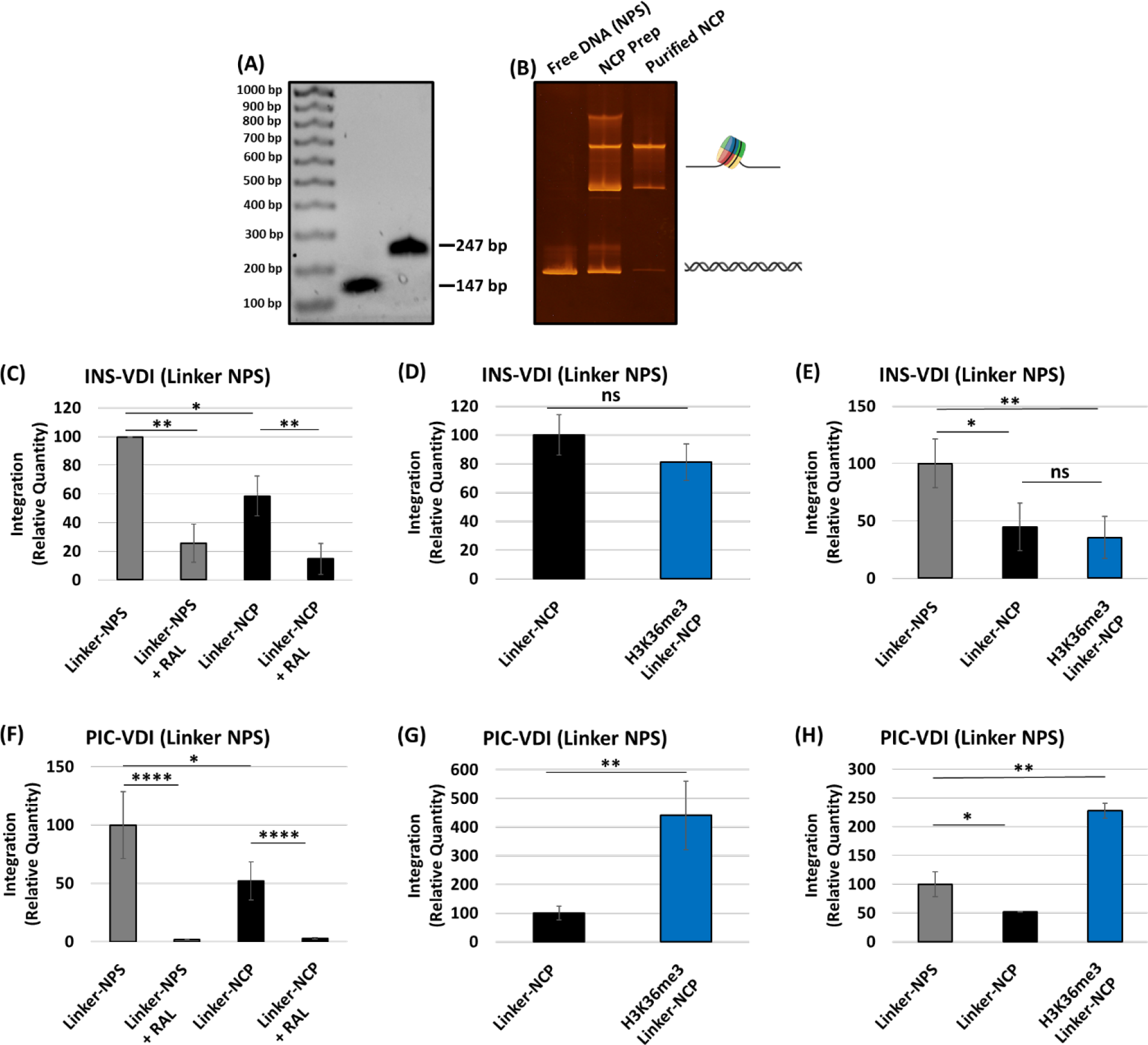
The NCP with linker DNA and the H3K36me3 is an optimal substrate for PIC-VDI. The nucleosome containing 50 bp of linker DNA flanking the Widom 601 NPS (linker-NPS) was assembled, resulting in an NCP with linker substrate (linker-NCP) that is 247 bp. (A) The 147 bp and 247 bp NPS DNA was analyzed on an 1.5% agarose gel and visualized by ethidium bromide staining. (B) Biochemical assembly of nucleosome with the 247 bp NPS was analyzed by EMSA. (C) The relative quantity of INS-VDI was measured with the linker-NPS, compared to the linker- NCP. The integrase inhibitor, RAL, was included as a control for VDI. (D) The INS-VDI with linker- NCP in reference to the H3K36me3 NCP was plotted. (E) The linker-NPS, linker-NCP and linker- H3K36me3-NCP substrates were compared for INS-VDI relative to the linker-NPS. (F) PIC-VDI was assessed with the linker-NPS and linker-NCP substrates. (G) The linker-NCP and linker- H3K36me3-NCP were compared for the relative PIC-VDI. (H) The PIC-VDI was measured comparing the linker-NPS, to the linker-NCP and the linker-H3K36me3-NCP. All the results are shown as relative integration quantity and represent the mean of at least three independent experiments. Error bars represent the SEM and * Represents P > 0.05, ** P = 0.01 to 0.05, *** P = 0.01 to 0.001, P = 0.001 to 0.0001, and **** P < 0.0001.

Furthermore, these data also indicate the PIC contains the necessary host factors to support such stimulation compared to the INS.

### LEDGF/p75 selectively stimulates PIC-VDI into specific substrates

Our results indicated that the nucleosome features (histone tail modification and DNA length) and the chromatin binding protein LEDGF/p75 directly regulate viral DNA integration. Particularly, the combination of the H3K36me3 and linker DNA within the NCP has an additive effect on PIC-VDI (Fig. 5H). This is in contrast to the PIC-VDI into non-linker NCP that showed a dramatic reduction in the presence of LEDGF/p75 (Fig. 3F). To better understand these paradoxical effects of LEDGF/p75, we measured PIC-VDI using the linker substrates in the presence of LEDGF/p75. We observed that with the H3K36me3-NCP (lacking linker DNA), LEDGF/p75 addition did not inhibit VDI, rather it resulted in a slight increase in PIC-VDI (Fig. 6A). Comparison of this data with the unmodified-NCP indicate that PIC-VDI is supported in the presence of H3K36me3. Moreover, the addition of LEDGF/p75 to the H3K36me3-nucleosome opposed the dramatic reduction in VDI observed with the unmodified-nucleosome (Fig. 6B). Further analysis revealed a 3-fold recovery of PIC-VDI with substrates containing the H3K36me3-NCP over the unmodified-NCP (Fig. 6C). Intriguingly, LEDGF/p75 addition to the unmodified linker NCP resulted in an enhanced PIC-VDI (Fig. 6D), an observation in clear contrast to the dose dependent reduction of PIC-VDI with the unmodified-NCP (Fig. 4D). Finally, we assessed PIC-VDI into the linker H3K36me3-NCP with LEDGF/p75 addition. To our surprise, these results showed no significant effect of LEDGF/p75 addition on PIC-VDI (Fig. 6E). Importantly, the raw integration values indicate that the integration quantity is relatively high (SI-Fig. 6C) suggesting that *in vitro* integration maybe occurring at an optimal level with the linker H3K36me3-NCP and exogenous LEDGF/p75. These observations provide further support that the presence of linker DNA and H3K36me3 in an NCP substrate is optimal for HIV-1 integration.

**Figure 6.**
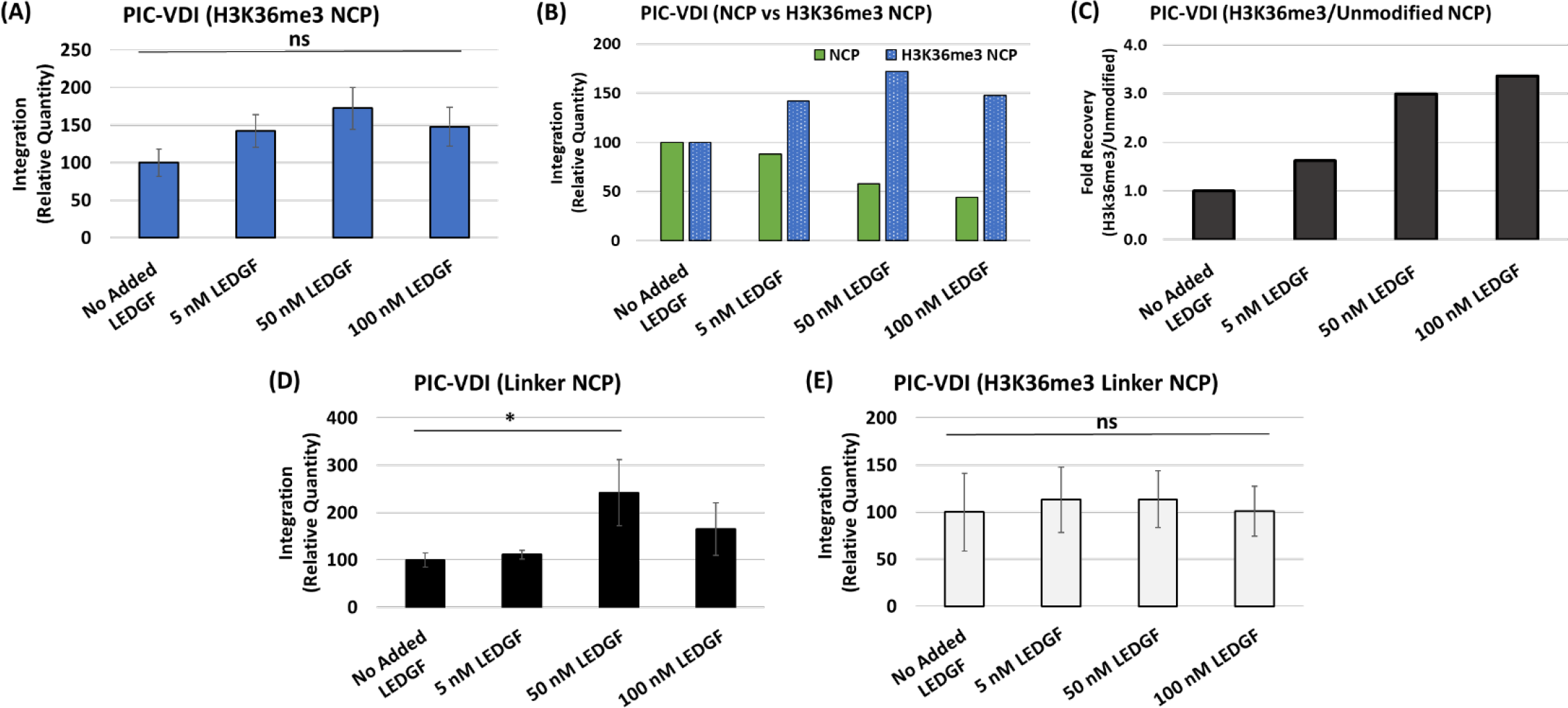
The H3K36me3-NCP and linker DNA support PIC-VDI in the presence of LEDGF/p75. (A) PIC-mediated integration was measured using H3K36me3-nucleosomes as targets in the presence of LEDGF/p75 [5, 50 and 100 nM]. Integration data are presented relative the PIC reactions without added LEDGF/p75. (B) Comparative analysis of the relative PIC-VDI with the NCP and H3K36me3-NCP in the presence of LEDGF/p75. (C) The fold change was calculated between the unmodified NCP and H3K36me3-NCP effect on PIC-VDI in the presence of LEDGF/p75. (D) PIC-VDI was measured with LEDGF/p75 addition to the linker-NCP substrate. (E) PIC-VDI with the linker-H3K36me3-NCP in the presence of LEDGF/p75 is plotted relative to the assay without LEDGF/p75 supplementation. The error bars represent the SEM of at least three independent experiments and * Represents P > 0.05, ** P = 0.01 to 0.05, *** P = 0.01 to 0.001, P = 0.001 to 0.0001, and **** P < 0.0001.

### The nucleosome core is preferentially targeted for integration when modified with H3K36me3

Our results identified that nucleosomes with a linker DNA and histone tail modification akin to *open chromatin* optimally enhanced PIC-VDI (Fig. 5H and SI-Fig. 6C). To further study the details of viral DNA integration into these preferred substrates, we deep sequenced integration reactions with the linker substrates ^79^. Duplicate integration reactions of the linker NPS, linker NCP and linker H3K36me3-NCP substrates were PCR amplified, concentrated, and spectrophotometrically analyzed. Amplicons were deep sequenced and we quantified reads that contained sequences of the HIV-1 5’ LTR adjoined to the linker sequence for the NPS-DNA, NCP, and H3K36me3-NCP targets. Analysis of the sequencing data identified integration junctions and revealed distinct integration patterns in each of the linker NPS, linker NCP, and linker H3K36me3-NCP substrates (SI-Fig. 8-10). With the linker NPS, integration junctions were identified throughout the length of the sequence with overrepresentation of viral DNA integration near the target DNA ends (Fig. 7A and SI-Fig. 8). Similarly, the integration junctions were also found in the unbound linker DNA regions of the linker NPC (Fig. 7B and SI-Fig. 9). Notably, the integration junctions within the linker H3K36me3-NCP were identified predominantly in the nucleosome core sequence, in striking contrast to the linker NPS and NCP (Fig. 7C, SI-Fig. 7 and SI-Fig. 10). The location of the integration sites within the linker H3K36me3- NCP are mapped near the entry/exit sites of the NCP (data not shown). Importantly, this is the location of the H3-tail protrusion between the nucleosomal DNA^39, 95, 98^, indicating that the HIV-1 PIC is drawn toward the H3K36me3 mark when engaged with a nucleosome during an integration event. To further understand these sequencing data, we quantified the integration site counts as a percentage of the total integration events. These analyses revealed that the linker NPS and linker NCP comprised ∼87 and ∼67 % of the junctions within the linker DNA, respectively (Fig. 7D). Distinctly, the linker H3K36me3-NCP contained over ∼90 % of the integration junctions adjoined in the nucleosome core sequence. Integration frequencies varied dramatically between the unmodified and H3K36me3-NCP, wherein the nucleosome core of the linker H3K36me3-NCP contained ∼95 % of the integration junctions (Fig. 7E, SI-Fig. 7B). However, the frequency of integration within the nucleosome core was only ∼15 % with the linker NPS-DNA and linker NCP lacking any histone modifications (Fig. 7E, SI-Fig. 7B). Collectively, these results indicate that H3K36me3 modification can regulate HIV-1 integration preference within a nucleosome.

**Figure 7.**
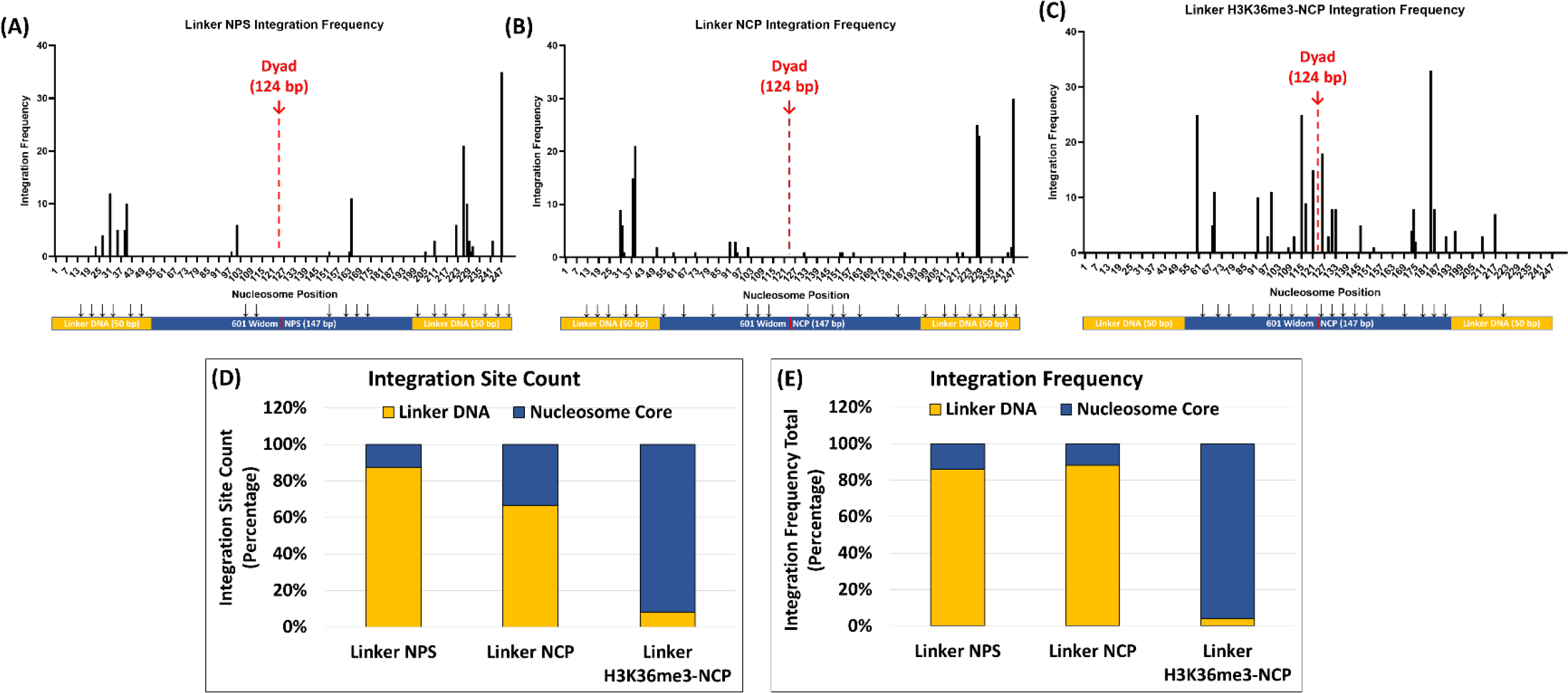
HIV-1 DNA integration is preferentially directed into the core of the H3K36me3 containing nucleosomes. To study HIV-1 integration preference, the DNA from the integration reactions were PCR amplified and subjected to next generation sequencing. The integration frequency within the linker substrates was determined by quantifying the integration junctions. The integration frequency is plotted as a histogram for (A) the linker-NPS, (B) linker-NCP, and linker-H3K36me3-NCP substrates. (D) After quantifying the integration sites within the linker- substrates, the percentage of sites within the linker sequences and nucleosome core sequence were plotted for comparative analysis. (E) Then the frequency of integration junctions at a particular site within the sequence was determined. The integration frequency in the linker sequences and the nucleosome core sequence was quantified as a percentage of the total integration sites.

**Figure 8.**
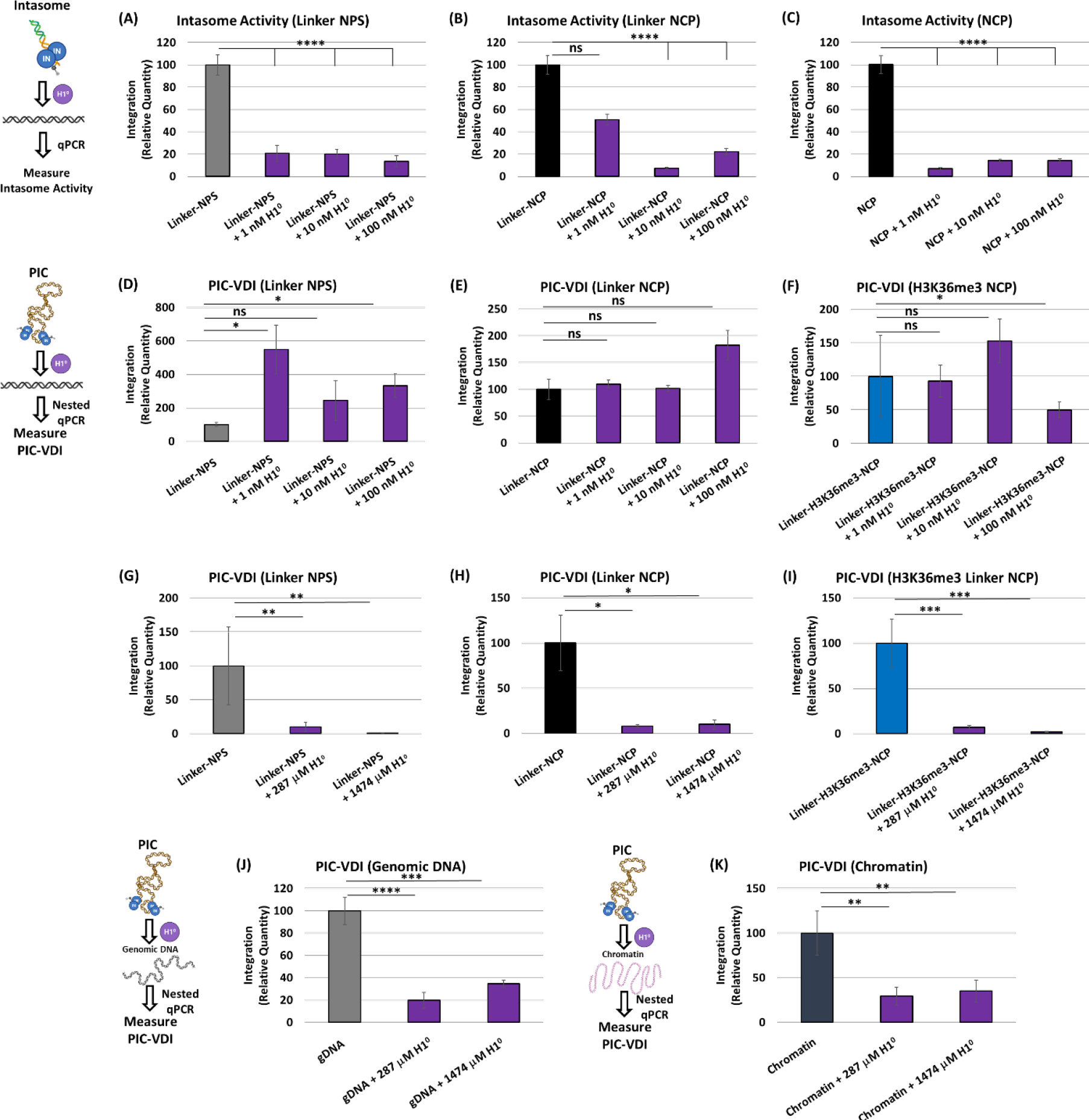
H1^0^ reduces HIV-1 DNA integration. To probe the effects of H1 on HIV-1 integration, first INS-VDI [25 nM] was tested with (A) the linker-NPS, (B) the linker-NCP or (C) the NCP substrate preincubated with recombinant H1^0^ [1, 10 or 100 nM]. The results are the average relative quantity in reference to the assay lacking H1^0^ addition. Then, PIC-VDI was measured with the linker-NPS, (E) the linker-NCP, (F) and the H3K36Cme3-linker-NCP substrates, that were preincubated with 1, 10 or 100 nM of H1^0^. (G-I) Next, either 287 or 1474 µM H1^0^ was added to the PIC-VDI measurements with the linker-NPS, the linker-NCP, and the H3K36Cme3-linker-NCP substrates. The µM concentrations of H1^0^ reflect amounts that are, respectively, 1:1 and 1:5 (w/w) stoichiometry of the substrate concentrations. The data shown is the relative quantity of the PIC integration relative to the assay lacking H1^0^ addition. The PIC-VDI was measured with (J) genomic DNA and (K) chromatin that were incubated on ice with 287 and 1474 µM of H1^0^. All data represent the mean of at least three independent experiments with the error bars representing SEM and * represents P > 0.05, ** P = 0.01 to 0.05, *** P = 0.01 to 0.001, P = 0.001 to 0.0001, and **** P < 0.0001.

**Figure 9.**
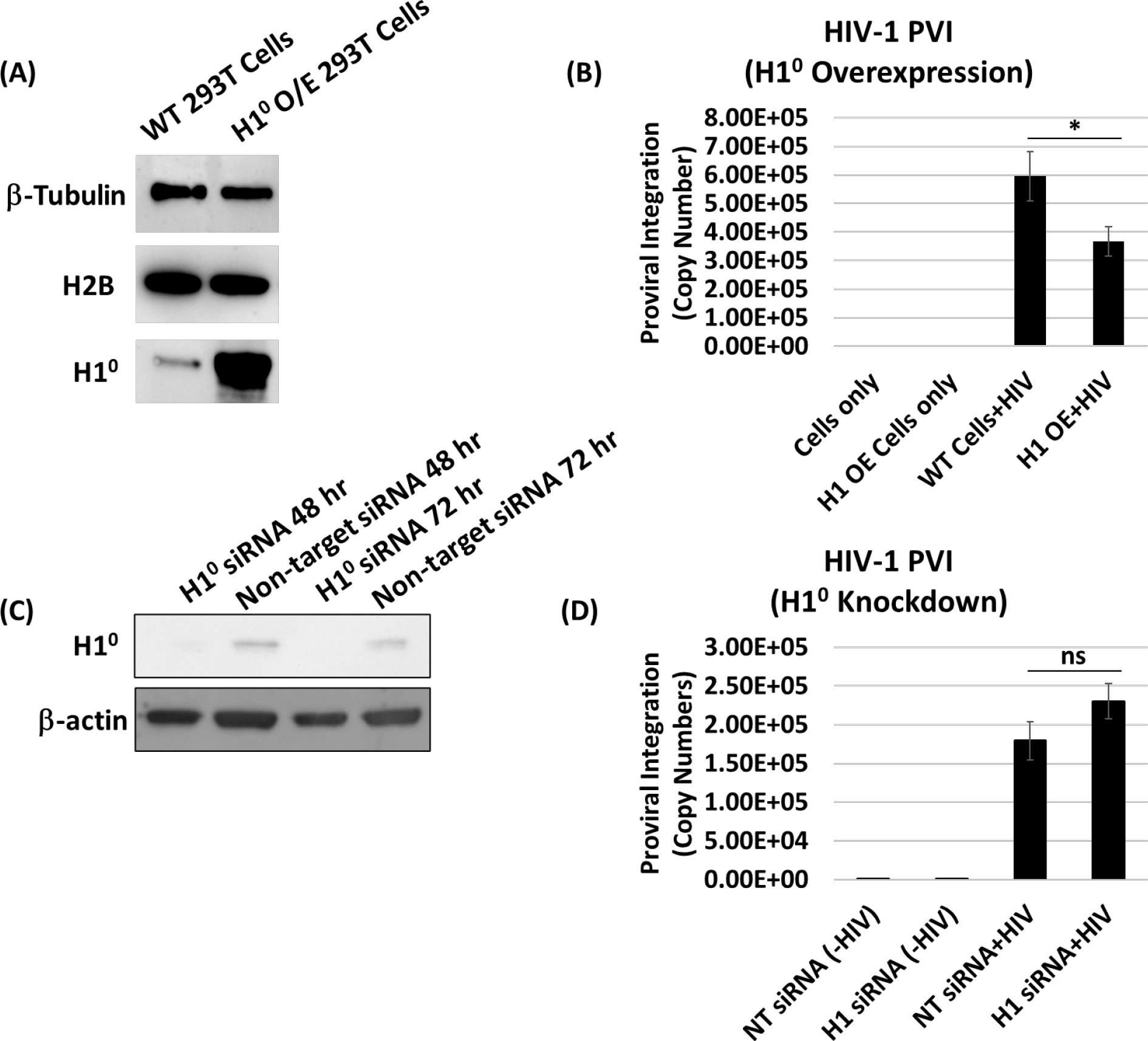
Histone H1 expression negatively regulates HIV-1 integration. (A) Linker histone H1^0^ was overexpressed in HEK 293T cells, and H1^0^ protein was probed by western blot analysis, and the same blot was re-probed for β-tubulin and histone protein H2B as loading controls. (B) Proviral integration in the HEK 293T cells overexpressing H1^0^. Cells were transfected with the pEV833 (GFP) expression construct for H1^0^, then were inoculated with VSVG-Δenv-HIV-1 (GFP) particles. 24 h post inoculation, HIV-1 proviral DNA integration (PVI) was quantified by nested Alu-PCR. (C) In TZM-bl cells, H1^0^ expression was depleted by siRNA targeting the H1-0 linker histone gene. Western blot analysis confirms the depletion of H1^0^ protein levels compared to scrambled siRNA control. The same blot was re-probed for β-actin. (D) The PVI of HIV-1 was measured in H1^0^ expressing and H1^0^ knockdown cells. All PVI data represent at least three independent experiments, and the error bars represent the SEM, * represents P > 0.05, ** P = 0.01 to 0.05, *** P = 0.01 to 0.001, P = 0.001 to 0.0001, and **** P < 0.0001.

### Chromatin compacting Histone H1 inhibits HIV-1 DNA integration

To further study the role of chromatin structure on HIV-1 integration, we next tested effects of H1 protein on HIV- 1 DNA integration. It is well-established that H1 reduces chromatin accessibility by facilitating chromatin compaction ^37, 100^ through the direct interaction with the linker DNA near the entry/exit location of the NCP and at the NCP dyad ^36, 101^. Therefore, we measured HIV-1 DNA integration by both PIC and INS in the presence of increasing concentrations of purified recombinant histone H1^0^ protein (0-100 nM). Our results revealed that INS-VDI was dramatically reduced in a dose-dependent manner with increasing concentration of H1^0^ protein (Fig. 8A-C). The inhibitory effect of H1 was universally observed with the linker NPS-DNA, linker NCP and the non-linker NCP. Surprisingly, addition of H1^0^ protein did not inhibit PIC-VDI at the concentrations of 0-100 nM (Fig. 8D-F). Rather with the linker substrates, nM concentrations of H1^0^ either slightly enhanced PIC- VDI with the linker NPS-DNA or had a minimal effect on PIC-VDI with both the unmodified and H3K36me3 linker NCP substrates (Fig. 8D-F). Then, we tested H1^0^ addition at saturating concentrations, 287 and 1474 µM, which correspond to a 1:1 or 1:5 molar ratio to the substrate concentrations, respectively. Interestingly, at these saturating concentrations, PIC-VDI was dramatically reduced into the NPS-DNA, NCP and H3K36Cme3 NCP substrates containing the linker DNA (Fig. 8G-I). Finally, we tested the effects of H1 on PIC-VDI using chromatin and de- chromatinized gDNA substrates. After preincubating the substrates with saturating concentrations of H1^0^, we observed significantly reduced PIC-VDI with the genomic DNA and chromatin substrates (Fig. 8J-K, SI-Fig. 13E-F). Notably, nM amounts of H1^0^ addition did not reduce PIC- VDI with the genomic DNA and chromatin substrates (SI-Fig. 12A-D), highlighting that specific stoichiometry between H1 and the chromatin is required for blocking accessibility of the PIC-associated viral DNA into chromosomal DNA. Collectively, these biochemical results provide further evidence of chromatin structure requirements for efficient HIV-1 DNA integration.

### Histone H1 regulates HIV-1 proviral integration

Our H1 protein addition results shows that H1^0^ can directly inhibit HIV-1 DNA integration. To test the physiological relevance of this biochemical observation, we probed whether H1 expression regulates HIV-1 DNA integration into human chromosomes in infected cells. To assess the effect of H1 on HIV-1 integration, we overexpressed H1^0^ in HEK293T cells prior to inoculation with psuedotyped HIV-1 virions. H1^0^ overexpression in these cells were confirmed by western blot analysis (Fig. 9A). The genomic DNA was extracted from the WT and H1^0^ overexpressing cells and the proviral integration (PVI) was measured by nested Alu-gag PCR method ^73, 74, 102^. Our results revealed that viral DNA integration was significantly reduced in the H1^0^ overexpressing cells relative to the control cells (Fig. 9B), suggesting a negative correlation between H1 expression and HIV-1 integration. We next carried out knocked-down studies using siRNA-based approach. TZM-bl cells transfected with H1-targeting siRNA showed dramatic reduction in H1 protein expression relative to the non-targeting control cells (Fig. 9C). Measurement of PVI in these cells showed that H1-depletion resulted in a slight increase in HIV-1 DNA integration levels (Fig. 9D). Collectively, these results provide evidence that H1 could play a role in blocking HIV-1 integration.

## Discussion

HIV-1 infection is dependent on the integration of the viral DNA into genic hotspots of host chromosomes ^2^. These hotspots contain actively transcribing genes that are epigenetically marked with histone modifications of *open chromatin* ^21, 25, 26, 29–31^. However, the exact role of viral and host factors that program the PIC to integrate the viral DNA into these hotspots remains largely unclear. Furthermore, establishing a direct and functional link between the markers of *open chromatin* and HIV-1 integration preference in cell-based and *in vitro* systems has been challenging ^51, 53, 54, 58, 103^. Therefore, we employed a biochemical approach involving the extraction of PICs from acutely infected cells and quantified viral DNA integration activity *in vitro* (SI-Fig. 1A-B) ^72–74^. We coupled PIC-VDI with in vitro assembled nucleosomes to define target substrate preference for HIV-1 DNA integration. Using this novel approach, we identified that the H3K36me3 histone modification-an epigenetic histone mark of active transcription, plays a role in HIV-1 DNA integration preference. Particularly, our results show that nucleosomes assembled with the H3K36me3 histone is a preferred target and HIV-1 DNA integration predominantly occurred within the DNA wrapped around the nucleosome core. The host-factor LEDGF/p75 stimulated HIV-1 DNA integration into the nucleosomes containing a linker DNA. Finally, the histone H1 protein that induces compaction of chromatin, dramatically reduced HIV-1 DNA integration into nucleosomes, naked DNA and chromatin substrates. Collectively, these studies establish a biochemical link between *open chromatin* structure and HIV-1 integration preference.

The molecular and biochemical mechanisms that drive HIV-1 DNA into specific regions of *open chromatin* are not clearly understood. This is in part due to the complexity of the structural landscape of the human chromatin coupled with an unclear understanding of the PIC function and composition. Chromosomes are oligomers of nucleosomes, which regulate access to the genomic DNA, through a number of intrinsic biochemical and structural mechanisms ^39–41^. Each nucleosome constitutes the NCP, linker DNA, and a linker histone ^34^. The NCP consists of ∼147 bps of DNA wrapped in left-handed super helical turns around a histone octamer ^35^. The histone octamer contains two copies each of the core histones H2A, H2B, H3, and H4 and the NCPs are connected by stretches of linker DNA ^75, 99^. Together, the NCP forms a disc-like structure that protects the DNA within the NCP from nucleases ^104^ and restricts the binding of transcription factors ^105, 106^. However, these stable nucleosomes must also allow access to the nucleosomal DNA for genomic functions such as DNA replication, RNA transcription and DNA damage repair ^107–109^. Evidently, this is achieved through the near meta-stable nature of the NCPs, driven by structural and conformation mechanisms involving: a) the intrinsic binding affinity of the histone octamers, b) post-translational modifications of the histone proteins and the DNA, c) the competitive or cooperative binding of other chromatin binding factors, and d) active translocation by ATP-dependent remodeling complexes. However, the mechanisms by which retroviruses counter the intrinsic structural and conformational barriers to access the nucleosomal DNA to insert the viral DNA is not fully understood. Nonetheless, *in vitro* studies of MLV PICs, together with purified MLV and HIV integrases (IN) show preference of the nucleosomal DNA relative to naked DNA as targets of integration ^43–46^. Furthermore, the curvature of nucleosomal DNA bending around the histone octamers or even the histone proteins themselves seem to impact viral DNA integration preference ^46, 47, 54, 103, 110^. Conversely, integration activity of recombinant HIV-1 IN enzyme can also be reduced with biochemically assembled nucleosomes and chromatin arrays are used as target substrates ^50, 53, 55^. Our results showing enhanced viral DNA integration into extracted chromatin-bound DNA relative to the de-chromatinized genomic DNA (Fig. 1D-E), provides the first biochemical evidence for the substrate preference of the HIV-1 PIC. These results are highly significant given that the PIC is solely responsible for carrying out HIV-1 DNA integration into the host chromosomes to establish life-long infection.

To understand the mechanism of chromatin preference for HIV-1 DNA integration, we adapted an *in vitro* reconstitution system to assemble nucleosomes with the Widom 601 nucleosome positioning sequence (Fig. 2A-D) ^77, 78^. This method generates compositionally uniform and precisely positioned nucleosomes compared to the cellular sources that contain highly diverse nucleosomes marked with an array of histone and DNA modifications ^40, 104, 111^. Most importantly, the biochemically assembled nucleosome is uniquely suited to test whether the NCP without any histone tail modification and the analogous NPS-DNA can serve as substrates for HIV-1 DNA integration. To our surprise, we observed higher viral DNA integration with the NPS- DNA rather than the NCP-bound DNA by both HIV-1 PICs and recombinant intasomes (INS) (Fig. 2E-F). Since, these NCPs are devoid of any histone tail modifications, our results suggest that the DNA-wrapped around naked nucleosomes lack the biochemical and/or structural determinants to support efficient HIV-1 DNA integration. Furthermore, the NPS-DNA (Widom 601) used in these experiments is optimized for binding to the NCP with high affinity ^76, 78, 80^. This could impose strong steric hindrance for the access and insertion of the viral DNA into NCP-bound DNA, whereas the naked NPS-DNA substrate possess no such biochemical or structural limitation. Notably, the previous studies showing enhanced integration into nucleosome-bound DNA were derived from disparate chromatinized substrates, such as chicken erythrocytes, yeast and SV40 mini-chromosomes, a mouse mammary tumor virus sequence mono-nucleosome, and a heterogeneously positioned chimeric di-nucleosome ^43–46^. Moreover, the histones used in subsequent nucleosome reconstitution studies were from a cellular source (HeLa cells or chicken erythrocytes) that most likely retained the native post-translational modifications that could influence viral DNA integration ^48–50, 55, 58, 79^. Therefore, our results suggest that the DNA wrapped within the NCPs lacking any histone tail modifications or linker DNA sequences is not a preferred substrate for HIV-1 DNA integration.

Chromatin structure is broadly categorized into *open* or euchromatin and *closed* or heterochromatin state ^75, 112^. Heterochromatin represents a highly condensed, gene-poor, and transcriptionally silent state, whereas euchromatin is less condensed, gene-rich, and more easily transcribed ^113^. Multiple features of chromatin, including histone modifications, DNA methylation, and small RNAs, are involved in higher order chromatin structure and thus, facilitate or prevent access to the nucleosomal DNA ^114, 115^. Notably, histone tail modifications characteristic of *open chromatin* are positively correlated with HIV-1 DNA integration preference ^21, 24–26, 28–30, 85, 89^ . Conversely, histone modifications that define DNA compaction into a heterochromatin state are negatively associated with HIV-1 integration. Particularly, the H3K36me3 modification is positively correlated with HIV-1 DNA integration across a number of published data sets ^21, 22, 25, 26, 29–32, 85, 116^. Although the underlying mechanism by which H3K36me3 enhances HIV-1 integration targeting is unclear, H3K36me3 is an abundant and highly conserved chromatin modification within the body of transcriptionally active genes ^117–119^. H3K36me3 also plays critical roles in the regulation of transcription elongation, DNA repair, prevention of cryptic start sites, pre-mRNA splicing and processing ^120–123^. Seemingly, H3K36me3 may play a role in tethering the PIC to the actively transcribing genes to facilitate HIV-1 DNA integration through the host factor LEDGF/p75 ^29, 30, 51, 98, 123, 124^. Accordingly, we observed higher PIC-mediated DNA integration into the nucleosomes containing H3K36me3 compared to the unmodified NCP (Fig. 3C and Fig. 5G). Quite surprisingly, the level of viral DNA integration into the H3K36me3-NCP was lower compared to the NPS-DNA (Fig. 3D and SI-Fig. 3B). Even though our results cannot fully explain the exact mechanism, the nucleosomes used in our study contain a single histone modification in contrast to the nucleosomes within native chromosomes (or natively sourced histones) that are decorated with a vast array of epigenetic histone and DNA modifications ^43–46, 48–50, 53, 55, 58, 79^. Interestingly, the presence of a linker DNA optimally stimulated PIC-mediated DNA integration into the linker H3K36me3-NCP (Fig. 5H and SI-Fig. 5F). The preference for the linker H3K36me3-NCP by the PICs was not observed with the HIV-1 intasome-mediated viral DNA integration (Fig. 3E-F and Fig. 5D-E), consistent with other studies that reported negligible effects by histone PTMs within chromatinized substrates on HIV-1 intasome activity ^51, 54, 58^. These results revealed that the recognition of H3K36me3 mark is PIC specific and is probably mediated by a viral and/host factor(s) other than the HIV-1 integrase enzyme. Interestingly, the HIV-1 PIC preference for integration with the linker H3K36me3-NCP aligns with studies suggesting HIV-1 IN targeted regions of chromatin arrays (assembled with native histones) with less nucleosome density as such will contain linker DNA ^34, 50, 55^. Taken together, these results suggest that H3K36me3 modification in the NCP and flanking linker DNA serve as biochemical determinants for efficient HIV-1 DNA integration.

During HIV-1 infection, a number of host factors are utilized to complete the early and late stages of the viral replication cycle ^125^. HIV-1 DNA integration into the host chromosomes by the PIC is dependent on host factors such as LEDGF/p75, INI1, CPSF6, and others ^89–91, 126^. The PIC-associated integrase procures LEDGF/p75 by interacting with the integrase binding domain (IBD) ^63, 92, 94^. Concomitantly, LEDGF/p75 specifically interacts with the H3K36me3-marked chromatin, and other methylated histones, through the PWWP domain and AT-hook motifs ^97, 98, 127^. It has been reported that LEDGF/p75 facilitates HIV-1 DNA integration through these binding interactions by tethering the PIC to actively transcribing genes ^49, 61, 63, 67, 94, 128^. Therefore, we hypothesized that the combination of LEDGF/p75 and the H3K36me3 would yield an additive effect on HIV-1 DNA integration in our *in vitro* assay. Accordingly, significant increase in HIV-1 DNA integration into both the NPC and NPS-DNA was observed by the intasomes in the presence of LEDGF/p75 (Fig. 4A-B). Surprisingly, LEDGF/p75 significantly reduced PIC-VDI with the NCP substrate and had minimal effect with the NPS-DNA (Fig. 4C-D). Notably, endogenous LEDGF/p75 is present in the extracted PICs (Fig. 4E), and addition of exogenous LEDGF/p75 blocked access to the NCP bound DNA (Fig. 4F-H). Thus, we predict that excessive LEDGF/p75 can sterically hinder access of the PIC-associated viral DNA to the NCP-bound target DNA. A lack of such block to the intasome-associated viral DNA, points to the involvement of additional factors and/or mechanisms specific to the PIC. Interestingly, LEDGF/p75 addition to PIC-mediated integration assays with the H3K36me3-NCP substrate relieved the steric block and allowed recovery of the PIC-mediated integration by three-fold (Fig. 6A-C). Furthermore, the presence of the flanking linker DNA that are exposed in *open chromatin*, resulted in the stimulation of PIC-mediated integration by LEDGF/p75 even with the unmodified NCP substrate (Fig. 6D). Collectively, these data implicate that structural and or conformational features of nucleosomes rendered by linker DNA and the H3K36me3 modification are essential for the host factor LEDGF/p75 to promote PIC-mediated viral DNA integration.

Our biochemical data shows that the H3K36me3-NCP containing linker DNA is the optimal substrate for PIC-mediated integration. However, our *in vitro* integration assay is not designed to identify the targets of integration into specific regions of the NCP. The NCP-bound DNA is highly contorted and bent into ∼1¾ left-handed super helical turns of about 80 bp/turn ^34, 35^. This bending is a consequence of the minor and major groove structures of the DNA where the minor grooves are preferentially oriented toward the histone octamer surface ^34, 78, 129, 130^. Notably, there is evidence that outward-facing DNA major grooves are favored target sites of retroviral integration ^45, 46^ and periodicity for integration occurs at the sites of the major grooves of the target DNA within the nucleosome. Even though, integrases lack DNA sequence specificity for integration targeting ^131, 132^, studies of mono-nucleosome particles ^45^ or mini-chromosomes ^43, 44^ have shown that the DNA wrapped around the nucleosome are preferentially targeted compared to nucleosome-free DNA. Therefore, to better understand the structural consequence of the H3K36me3-NCP on viral DNA targeting, we carried out deep sequencing of the viral DNA integrants from PIC-VDI assays with the linker NPS-DNA, linker NCP, and the linker H3K36me3-NCP. PIC-mediated viral DNA integration within the linker NPS preferably occurred throughout the DNA sequence and primarily in the linker sequence positions 1–50 bp and 197 – 247 bp (Fig. 7A, SI-Fig. 7 and SI-Fig. 8). Similarly, integration within the linker NCP occurred most frequently in the linker DNA and a slight uptick in the integration sites throughout the nucleosome core positioning sequence, base pairs 50 – 197 (Fig. 7B, SI-Fig. 7 and SI-Fig.9). Remarkably, the integration into the nucleosome core DNA increased within the linker H3K36me3-NCP. Both the integration count and frequency occurred overwhelmingly in the nucleosome core sequence of the H3K36me3-NCP with linker DNA (Fig. 7C, SI-Fig. 7 and SI-Fig. 10). It should be noted that these integration site analyses were from sequencing reads of <500, due to the technical challenges of shorter DNA substrates. Therefore, we are cautious about the statistical and physiological significance of these observations. Nevertheless, qualitative analysis of the integration sites within the linker H3K36me3-NCP indicate that the integrants abutted within the super helical location (SHL) -1, - 7, +7 and +1, which corresponds to the location where the H3-tail protrudes through the nucleosomal DNA ^39, 86, 98^. This striking observation suggests that the H3K36me3 mark can perform a functional role in directing the HIV-1 PIC-associated DNA into specific regions of the nucleosomal-DNA.

We provided biochemical evidence that nucleosome structure characteristic of *open chromatin* enhances HIV-1 integration. Interestingly, *open chromatin* represents a minority of the human genomic DNA, whereas the *closed chromatin* or heterochromatin represents up to 80 percent of the genomic DNA ^112, 133^. However, it is unclear why HIV-1 integration is disfavored in these vast DNA regions of the human chromosomes ^21, 26, 28, 29, 32^. To interrogate the role of closed chromatin on HIV-1 integration, we probed the effects of the linker histone H1^0^ due to its relatively high abundance among H1 variants ^134^ and well-characterized ability to condense chromatin fibers and form rigid, stable *chromatosome* structures ^36, 101^. As expected H1^0^ addition protected isolated chromatin from micrococcal nuclease digestion (SI-Fig. 13A-D) and at non-saturating amounts, inhibited HIV-1 intasome-mediated viral DNA integration (Fig. 8A-C). This is consistent with the published studies with HIV-1 integrase and H1-mediated condensation of recombinant nucleosome arrays ^48, 100, 135^. Surprisingly, non-saturating amounts of H1^0^ failed to inhibit PIC- mediated viral DNA integration into the linker substrates (Fig. 8D-F and SI-Fig. 11A-C). However, at stoichiometric saturating levels relative to the DNA substrate, PIC-mediated viral DNA integration was significantly reduced with all the linker substrates (Fig. 8 G-I and SI-Fig. 11D-F), the genomic DNA and the chromatin (Fig. 8 J-K and SI-Fig. 13E-F). Our data indicates that chromatosome formation is essential for the H1-mediated inhibition of PIC-VDI (SI-Fig. G-H). Finally, proviral DNA integration was also reduced in H1^0^ O/E cells and conversely was enhanced in H1^0^ depleted cells albeit to a smaller degree (Fig. 9B and 9D). The moderate effect by the reduction the H1^0^ on HIV-1 integration in the cell-based assays most likely represents the functional redundancy of the H1 variants, some of which could compensate the inhibitory effect of the single variant depletion ^136–138^. Notwithstanding, our data suggests that the linker histone H1 can negatively regulate HIV-1 DNA integration most likely through chromatin compaction.

In conclusion, our study provides biochemical evidence and mechanistic insights into HIV- 1 integration preference into *open chromatin*. For the first time we defined a direct role of the H3K36me3 epigenetic mark in HIV-1 integration preference and identified the optimal substrate for HIV-1 PIC-mediated integration. Finally, our study demonstrates the importance of biochemical studies of the PIC to achieve a clear understanding of the mechanism of retroviral DNA integration.

## Acknowledgements

This work was supported by the National Institutes of Health grants R01 AI136740, R01 DA 042348, R56 AI122960, R24 DA036420, R25AI1647610, R01AI162694, and U54 MD007586 to C.D., and the support from the Research Centers in Minority Institutions (RCMI) grant U54MD007586 to J.P, as well as the training grants R25 GM059994 and F31 AI150488 in support of N.S. This work is also supported in part by the Meharry Translational Research Center (MeTRC) grant U54MD007593 and Tennessee CFAR grant P30 AI110527 from the National Institutes of Health to C.D. M.L. and R.C. are supported by the intramural research program of the National Institutes of Health. We also acknowledge Jarrod Johnson, University of Utah for the technical support with the sequencing analysis. We declare that we have no conflict of interest with the content of this article.

## Materials and Methods

### Cell culture

HEK293T and SupT1 cell lines were obtained from the American Type Culture Collection (Manassas, VA). The TZM-bl reporter cell line was obtained from John C. Kappes, Xiaoyun Wu, and Tranzyme, Inc., through the NIH AIDS Reagent Program, Division of AIDS, NIAID, NIH. HEK293T and TZM-bl cells were cultured in Dulbecco modified Eagle medium (DMEM) supplemented with 10% heat-inactivated fetal bovine serum (FBS), 2 mM glutamine, 1,000 U/ml penicillin, and 100 mg/ml streptomycin. SupT1 cells were cultured in RPMI 1640 medium supplemented with 10% heat inactivated FBS, 2 mM glutamine, 1,000 U/ml penicillin, and 100 mg/ml streptomycin. All cells were cultured at 37°C with 5% CO2.

### Virus stocks

High titer virus stocks were generated by transient transfection of HEK293T cells with the HIV-1 plasmid construct pNL4-3 (NIH AIDS Reagents) by with polyethyleneimine (PEI). Briefly, 3 X 10^6^ cells were seeded per 10-cm culture dish and cultured overnight. The following day, cells in each dish were transfected using PEI and 10-15 µg of pNL4-3 DNA. At 16 hours post-transfection, the cells were washed with phosphate buffered saline (1X PBS), replenished with 6 ml of DMEM. After 36 hours, the virus-containing culture supernatant of transfected cells were harvested, cleared of debris by low-speed centrifugation, filtered through 0.45-mM filters, and DNase I treated (Calbiochem; 20 µg/ml of supernatant) in the presence of 10 mM magnesium chloride (MgCl2) for 1 hour at 37°C. The virus infectivity was determined by using TZM-bl indicator cells, as previously described ^139^.

### Chromatin isolation

The chromatin isolation protocol is adapted from a published protocol ^140^. HEK293T cells were cultured to near confluency, then harvested and washed once with ice cold 1X PBS. Cells were manually lysed on ice with a Dounce homogenizer (Wheaton) in *native lysis buffer* (NLB: 5 mM MgCl2, 10 mM KCl, 20 mM HEPES pH 7.5, 250 mM sucrose, 0.5 mM dithiothreitol (DTT), 0.5 mM phenylmethanesulfonylfluoride (PMSF)) for 20 to 30 strokes. The lysate was washed and homogenized with NLB twice by centrifugation at 4°C on 3,000x g for 15 minutes. After manual lysis with NLB, the pellet was washed with modified buffer B (MBB: 5 mM MgCl2, 10 mM KCl, 20 mM HEPES pH 7.5, 0.2 M ethylene glycol-bis(2-aminoethylether)- N, N, N’, N’-tetra acetic acid (EGTA), 0.5 mM DTT, 0.5 mM PMSF, and protease inhibitor cocktail (Promega, Madison, WI, USA)). The resulting pellet was resuspended in 2 volumes of MBB, in reference to the pellet volume, and then an equivalent volume of MBB/0.6 M KCl/ 10% glycerol was added in a dropwise manner. The lysate was incubated at 4°C for 10 minutes with consistent rocking. After incubation, the nuclear fraction was pelleted at a pre-cooled (4°C) centrifuge at maximum speed. The pelleted nuclear fraction was resuspended in 20 times volume of medium salt buffer (MSB: 20 mM HEPES, pH 7.5, 0.4 M NaCl, 1 mM ethylenediaminetetraacetic (EDTA), 5% glycerol (v/v), 0.5 mM DTT and 0.5 mM PMSF) and manually homogenized. The nuclear pellet was centrifuged at 4°C for 10 minutes at 11,000x g, then resuspended in 4 pellet volumes of high salt buffer (HSB: 20 mM HEPES, pH 7.5, 0.65 M NaCl, 1 mM EDTA, 0.34 M sucrose, 0.5 mM DTT and 0.5 mM PMSF).

Oligonucleosomes were released from the nuclear debris by manual homogenization on ice, and fully separated from the debris by a maximum speed centrifugation at 4°C to collect the chromatin containing supernatant. The chromatin was dialyzed for 16 hours against low salt buffer (LSB: 20 mM HEPES, pH 7.5, 0.1 M NaCl, 1 mM EDTA, 0.5 mM DTT and 0.5 mM PMSF). The dialyzed chromatin was tested for nucleosome protection and the presence of histone proteins.

### Immunoblotting

To detect proteins by immunoblot, samples were prepared in Radioimmunoprecipitation assay (RIPA) buffer (Santa Cruz Biotechnologies Inc., Dallas, TX, USA) or the NLB buffer supplemented with protease inhibitor cocktail and 10 μg/µL PMSF (Sigma-228 Aldrich, Saint Louis, MO, USA) according to the standardized protocol. Protein concentrations were determined using BCA (Bicinchoninic Acid) Protein Assay reagent (Thermo-Fisher Scientific, Waltham, MA, USA) by the manufacturer’s specifications. Equivalent amounts of protein from cellular lysates or fractions were electrophoresed on pre-casted gels (Bio-Rad Laboratories, Hercules, CA, USA) or in-house prepared 15% polyacrylamide gels and electrophoretically transferred to nitrocellulose membranes using the Trans-Blot SD Semi-Dry Transfer Cell (Bio-Rad Laboratories). The membranes were incubated in blocking buffer [5% (w/v) nonfat milk in Tris-buffered saline containing 0.05% Tween 20 (TBST); pH 8.0]. After blocking the membranes were then probed with the indicated antibody diluted in blocking buffer; Abcam, Boston, MA, USA: H2A (ab18255), H2B (ab1790), H3 (ab1791), H4 (ab7311), H1^0^ (ab218417); Active Motif, Carlsbad, CA, USA: H3K36me3 (ab_2615073); Bethyl Laboratories, Inc., Montgomery, TX, USA:: LEDGF/P75 (A300- 848A); Cell Signaling Technology: p53 (2527S); Proteintech, Rosemont, IL, USA: GAPDH (60004-1), and subsequently with secondary antibody conjugated to horseradish peroxidase [anti-rabbit 1:10,000 (v/v); anti-mouse 1:10,000 (v/v)]. The membranes were washed with 1X TBST buffer at least 3 times for 15 minutes each wash, and immunocomplexes were detected by clarity enhanced chemiluminescence (ECL) method (Bio-Rad Laboratories).

### Isolation of preintegration complexes (PIC) from HIV-1 infected SupT-1 cells

HIV-1 PICs were isolated from acutely HIV-1 infected T cells as described by Balasubramaniam and colleagues ^71–74^. Briefly, 8 X 10^7^ of SupT1 cells were spinoculated (480 X *g*) with DNase I-treated wild type virions for 2 hours at 25°C, and then cultured for 5 hours at 37°C. The infected cells were then harvested by centrifugation (300 X *g*) for 10 minutes at room temperature. The cell pellet was washed twice with 2 ml of buffer K^-/-^ (20 mM HEPES, [pH 7.5], 150 mM potassium chloride (KCl), 5 mM MgCl2). Subsequently, the cell pellet was gently resuspended in 2 ml of ice-cold buffer K^+/+^ (20 mM HEPES, [pH 7.5], 150 mM KCl, 5 mM MgCl2, 1 mM dithiothreitol [DTT], 20 mg/ml aprotinin, 0.025% [wt/vol] digitonin). The cell suspension was transferred to a chilled 2 ml microcentrifuge tube and incubated on a rocking platform (60 to 80 rocking motions/ minute) for 10 minutes at room temperature. The cell lysate was centrifuged (1,500 X g) for 4 minutes at 4°C to separate the cytoplasmic and nuclear fractions. The supernatant (cytoplasmic fraction) was transferred to a fresh 2 ml microcentrifuge tube and centrifuged again (16,000 X g) for 1 minute at 4°C to clear residual nuclear debris. The resulting cytoplasmic faction was treated with RNase A (Invitrogen, Waltham, MA, USA) at a final concentration of 20 mg/ml and incubated for 30 minutes at room temperature. Finally, sucrose (60% [wt/vol]) was added to a final concentration of 7% and the contents were thoroughly mixed. Aliquots of the cytoplasmic fraction were flash-frozen in liquid nitrogen and stored at -80°C for long-term storage. The cytoplasmic fraction aliquots were used as the source of HIV-1 PICs in assays.

### Assay for measuring PIC-viral DNA integration (VDI)

The *in vitro* integration assays were performed using a modified version of a protocol detailed by Balasubramaniam and colleagues ^72–74^. The *in vitro* integration reaction was carried out by mixing 50 µl of PIC with 300 ng of the indicated target DNA, then allowing the mixture to incubate at 37°C for 45 minutes. The integration reaction was stopped by adding SDS, EDTA, and proteinase K to a final concentration of 0.5%, 8 mM, and 0.5 mg/ml, respectively, followed by an overnight incubation at 55°C. The deproteinized reaction was brought to 200 µl and mixed with an equal volume of phenol [pH 8.0] and thoroughly mixed by vortexing, and centrifuged (17,000 X g) for 2 minutes at room temperature. The aqueous phase is extracted once with an equal volume of phenol-chloroform (1:1) mixture, followed by an equal volume of chloroform. The DNA was precipitated by adding 2.5 volumes of 100% ice-cold ethanol in the presence of sodium acetate (0.3 M, final concentration) and the coprecipitant glycogen (25 to 100 mg, final concentration), followed by a minimum incubation for 2 hours at -80°C. The sample was centrifuged at maximum speed for 30 minutes at 4°C, and the resultant DNA pellet was washed once with 70% ethanol using maximum speed centrifugation for 15 minutes at 4°C. The precipitated DNA was air dried at room temperature, resuspend in 50 µl of nuclease free water, and used as the template DNA for the nested qPCR to measure PIC-mediated viral DNA integration.

The nested PCR consists of two rounds of PCR to amplify the junction between the viral DNA and target DNA, followed by a quantitative PCR (qPCR) specific to the viral DNA. A first-round standard PCR, designed to amplify only the integrated virus-target DNA junctions, was performed in a final volume of 50 µl containing 5 µl of purified DNA product from the integration reaction, 500 nM concentration of each primer against the indicated target DNA (Table S2) and the viral long terminal repeat (LTR) [5’-GTGCGCGCTTCAGCAAG- 3’], 1X Bestaq PCR buffer (Applied Biological Materials Inc., Richmond, BC, Canada), dNTP nucleotide mix containing 200 mM concentrations of each nucleotide (Promega, Madison, WI, USA), and 1.25 U of Bestaq DNA Polymerase (Applied Biological Materials Inc.) with the following thermocycling conditions: 95°C for 5 minutes, followed by 25 cycles at 95°C for 30 seconds, 55°C for 30 seconds, and 72°C for 30 seconds (for the Widom 601 NPS DNA) or 2 minutes (for chromatin or genomic DNA), and a final extension at 72°C for 10 minutes. The second-round qPCR designed to amplify on the viral LTR-specific region contained one-tenth (5 µL) the volume from the first-round PCR product as the template DNA, 1X iTaq Universal Probe Supermix (Bio-Rad Laboratories), 300 nM concentrations each of the viral LTR-specific primers that target the R region (5’- TCTGGCTAACTAGGGAACCCA-3’) and the U region (5’-CTGACTAAAAGGGTCTGAGG-3 ’), and a 100 nM concentration of TaqMan probe (5’-6-carboxyfluorescein[FAM]- TCAGCATTATCAGAAGGAGCCACC-6-carboxytetramethylrhodamine [TAMRA]-3’). The qPCR run included an initial incubation at 95°C for 3 minutes, followed by 39 cycles of amplification and acquisition at 94°C for 15 seconds and 58°C for 30 seconds, and a final incubation at 72°C for 30 seconds. During qPCR of the samples, a standard curve was generated under the same conditions in parallel using 10-fold serial dilutions of known copy numbers (10^0^ to 10^8^) of the HIV-1 molecular clone plasmid. Data were analyzed using CFX Manager software (Bio-Rad Laboratories), and integrated viral DNA copy numbers were determined by plotting the qPCR data against the standard curve.

### Nucleosome positioning sequence DNA preparation

The DNA molecules used for nucleosome assembly were generated by PCR from a pUC19 clone containing the Widom 601 nucleosome positioning sequence (NPS). DNA products from a 48-well PCR plate (50 - 100 µL reaction per well) were pooled and injected into an anion exchange column (MonoQ-GL, GE Healthcare Life Sciences, Marlborough, MA, USA) for purification. After extensive washing with 0.5X tris-EDTA (TE) buffer, the DNA was eluted with a 0 – 1.0 M NaCl linear gradient in 0.5X TE buffer, pH 7.5. The fractions containing the PCR product was pooled and concentrated by a minimum cut-off of 10 MWCO spin column (Amico Ultra, Millipore Sigma, Burlington, MA, USA) in 0.5X TE.

### Nucleosome Reconstitution

Nucleosomes were made as previously described ^77^. DNA was mixed with histone octamer in a molar ratio of 1.25:1 in 0.5X TE pH 8, 2 M NaCl, and 1 mM benzamidine hydrochloride and dialyzed via double dialysis into 0.5X TE pH 8 and 1 mM benzamidine hydrochloride. Reconstituted nucleosomes were separated from free DNA using ultracentrifugation with a 5 - 30% w/v sucrose gradient spun at 41,000 rpm for 22 hours. Gradients were fractioned and ran on a 0.3X TBE, 5% acrylamide gel at 300 V for 1 hour to determine nucleosome containing fractions. Selected fractions were concentrated, and buffer exchanged into 0.5X TE pH 8.0 using centrifugal filters and final concentrated nucleosomes were run on acrylamide gels as before to verify purity.

### Deposition of methyl-lysine analog on histone H3

Human H3.2 (C110A, K36C) was created by site-directed mutagenesis using a Stratagene QuickChange XL kit. The mutant histone was then expressed in *E. coli* BL21(DE3) pLysS cells and purified as previously described ^77^. The methyl-lysine analog was then deposited onto H3.2(C110A, K36C) based on a previously reported protocol ^87, 141^. Initially, 5 mg of the mutant histone was suspended in 980 µL of alkylation buffer (1 M HEPES pH 7.8, 4 M guanidine–HCl, and 10 mM D/L-Methionine) and allowed to unfold for 1 hour. Then histones were reduced with 6.66 mM dithiothreitol for 1 hour at 37°C. After reducing, 100 mg of (2-bromoethyl) trimethylammonium bromide was added to the histones. The reaction was then covered and stirred for 5 hours at 50°C. The reaction was quenched with 50 µL of 14.3 M 2-mercaptoethanol. Histones were then desalted using a PD 10 desalting column and dried via vacuum concentration. Labeling efficiency was confirmed by MALDI-TOF mass spectrometry.

### HIV-1 Intasome (INS) preparation and assembly

The HIV-1 intasomes used in our study were prepared by using published methods ^81, 82^. Briefly, a double-stranded preprocessed viral DNA substrate was used as the donor DNA (Integrated DNA Technologies, Coralville, IA, USA). A 25-base oligonucleotide (5’- AGCGTGGGCGGGAAAATCTCTAGCA-3’) was synthesized with a complementary 27-base oligonucleotide (5’-ACTGCTAGAGATTTTCCCGCCCACGCT-3’) to provide the 3’-OH recessed end for the preprocessed intasomes. The recombinant Sso7d-integrase (Sso7d-IN) was expressed and purified in Escherichia coli BL21 (DE3), and the cells were lysed in buffer containing 20mM HEPES (pH 7.5), 10% glycerol, 2mM β-mercaptoethanol (β-ME), 20mM imidazole, and 1 M NaCl. The protein was purified by nickel affinity chromatography and subsequently by gel filtration on a Hi load 26/60 Superdex-200 column (GE Healthcare, Milwaukee, WI, USA) equilibrated with 20mM HEPES (pH 7.5), 10% glycerol, 2mM DTT, and 500mM NaCl. The protein was concentrated using an Amicon centrifugal concentrator (EMD Millipore, Burlington, MA, USA) and then used for intasome assembly or flash frozen in liquid nitrogen and stored at -80°C. Intasomes were assembled by incubating 3 mM HIV-1 integrase and 1 mM 3’-processed or unprocessed viral DNA substrate in 20mM HEPES (pH 7.5), 20% glycerol, 5mM 2-mercaptoethanol, 5mM CaCl2, 2mM ZnCl2, 100mM NaCl, and 50mM 3- (benzyldimethylammonio) propanesulfonate (NDSB-256) at 30°C for 1 h. Assembled intasomes were purified as described previously ^81, 82^, aliquoted, flash frozen in liquid nitrogen, and stored at -80°C.

### Assay to measure HIV-1 Intasome-VDI

Intasome viral DNA integration (INS-VDI) assays were assembled on ice with 25 nM INS in 20 mM HEPES pH 7.5, 20% glycerol, 10 mM DTT, 5 mM MgCl2, 4 mM ZnCl2 and 100 mM NaCl in a 20 µl reaction volume. *In vitro* integration reaction with intasomes were carried out with 300 ng of target DNA, the incubated at 37°C for 1 hour. The reaction was terminated with 0.5% SDS and 8 mM EDTA, together with 0.5 mg/ml protease K and deproteinated for 1 hour at 55°C. The DNA was recovered by ethanol precipitation, as described above, and resuspend in the original reaction volume of nuclease-free dH2O. The recovered DNA was used as the template DNA for a SYBR green-based quantitative PCR to amplify the junctions between viral DNA substrate and the target DNA for the indicated assay. The qPCR reaction was carried out in a 20 µl volume, containing 1 to 2 µl (5 - 10 ng) of purified DNA product from the INS-VDI assay, 300 nM concentration of each primer targeting the viral DNA substrate (5’- AGCGTGGGCGGGAAAATCTC-3’) and the indicated target DNA (table 1) and 1X i*Taq* Universal SYBR Green Supermix (Bio-Rad Laboratories). The qPCR cycling conditions for quantifying INS activity included an initial incubation at 95°C for 3 minutes, followed by 39 cycles of amplification and acquisition at 94°C for 15 seconds, 55°C for 30 seconds and 72°C for 30 seconds. The thermal profile for melting-curve analysis was obtained by holding the reaction at 65°C for 30 seconds, followed by a linear ramp in temperature from 65 to 95°C at a ramp rate of 0.5°C/second and acquisition at 0.5°C intervals. The Maestro CFX program was used for data analysis.

### FRET Titrations

FRET efficiency measurements were carried out with a Horiba Scientific Fluoromax 4 Spectrofluorometer. Samples were excited at 510 and 610 nm and the photoluminescence spectra were measured from 530 to 750 nm and 630 to 750 nm for donor and acceptor excitations, respectively. FRET efficiencies from measured spectra were computed through the (ratio) A method ^142^ using an in-house Matlab program. FRET efficiencies were then normalized as change in FRET and fit to a binding isotherm using GraphPad 9.1.2 to determine S1/2’s, the concentration at which 50% of the nucleosome is bound by LEDGF/p75.

Titrations used a 60 µL reaction volume with 1 nM nucleosomes mixed into increasing concentrations of GAL4. Each titration was repeated in the presence or absence of binding saturation concentrations of LEDGF (500 nM). The conditions in each reaction were 75 mM NaCl, 10 mM Tris-HCl pH 8, 0.25% Tween20, and 10% glycerol. Titrations were repeated in triplicate to acquire standard deviation errors.

### Fluorescence polarization

Fluorescence polarization measurements were acquired with a Tecan infinite M1000Pro plate reader by exciting samples at 470 nm and then measuring polarized emission at 519 nm with 5 nm excitation and emission bandwidths. Fluorescence polarization of each sample was calculated from:

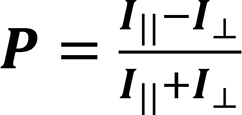

where *I*_||_ and *I*_⟂_ is the emission polarized parallel and perpendicular to the polarized excitation light, respectively. The data was then fit to a binding isotherm using GraphPad 9.1.2 to determine S1/2’s.

Titrations used a 30 µL reaction volume with 5 nM nucleosomes mixed into increasing concentrations of LEDGF. The conditions in each reaction were 75 mM NaCl, 15 mM Tris-HCl pH 7.5, 0.00625% Tween 20, and 3 mM dithiothreitol. The samples were loaded into a Corning round bottom polystyrene plate and allowed to incubate at 4°C for 30 mins before measurements were taken. Titrations were repeated in triplicate to acquire standard deviation errors.

### Proviral Integration (PVI) Assay

Infection experiments were carried out by inoculating cells with psuedotyped HIV-1 particles (ref). HEK293T or TZM-bl (3.0 x 10^5^ cells per well) were seeded in 6-well plates, cultured overnight, and then washed with fresh medium. Virus stocks prepared in DMEM containing Polybrene (6 µg/ml) and either 1 µM RAL, or 5 µM EFV were indicated were added to the cells. The cells were then cultured for 24 to 48 hours at 37°C with 5% CO2 and harvested for further use. Total DNA from the infected and uninfected control cells was isolated using a Quick-DNA miniprep kit according to the manufacturer-recommended protocol (Zymo Research, Irvine, CA, USA). To measure HIV-1 PVI, a nested PCR method was used that consisted of a first-round endpoint PCR with primers designed to amplify only the integration junctions between human *Alu* repeats and HIV-1 viral DNA, followed by a second-round of qPCR with primers designed to specifically amplify only the viral LTR from the first-round PCR products (ref). The first-round PCR contained 100 ng of total DNA, 1X Bestaq PCR buffer (Applied Biological Materials Inc.), dNTP mix containing 200 mM concentrations of each nucleotide (Promega, Madison, WI, USA), 500 nM primers targeting *Alu* repeat sequence (5’-GCCTCCCAAAGTGCTGGGATTACAG-3’) and the HIV-1 Gag sequence (5’-GTTCCTGCTATGTCACTTCC-3’), and 1.25 U of Bestaq DNA polymerase in a 50-µl final volume. The first-round PCR conditions consisted of the following cycles: 95° C for 5 minutes, followed by 25 cycles of amplification at 94°C for 30 s, 50°C for 30 s, and 72°C for 4 min, and a final incubation at 72°C for 10 min. The second-round PCR reaction consisted of one-tenth (5 µL) of the product from the first-round PCR as the DNA template, 1X i*Taq* universal probe Supermix (Bio-Laboratories), 300 nM each of the viral Gag specific primers (F2: 5’-TCAGCCCAGAAGTAATAC-3’) and (R2: 5’-CACTGGATGCAATCTATC-3’), and 100 nM of the TaqMan probe (5’-6-[FAM]- TCAGCATTATCAGAAGGAGCCACC-6-[TAMRA]-3’). The qPCR program consisted of the following amplification conditions: 95° C for 3 min, followed by 39 cycles of amplification and acquisition at 94°C for 15 s, 58°C for 30 s, and 72°C for 30 s. Concurrent with the qPCR of the samples, a standard curve was generated in parallel and under the same conditions using a 10-fold serial dilution of known copy numbers (1 X 10^0^ to 1 X 10^8^) of the HIV-1 molecular clone plasmid (NL4-3). The qPCR experiments were performed in triplicates, and the data were analyzed using CFX Maestro software (Bio-Rad Laboratories). The integrated viral DNA copy numbers were calculated by plotting the qPCR data against the standard curve.

### Sequencing Analysis of the integration junctions

The first-round end point PCR product of the PIC-VDI assay was subjected to a second end point PCR amplification with the same primers to increase the yield of DNA. The second endpoint PCR amplification was subjected to a PCR DNA purification kit (Zymo Research) and the DNA was submitted for sequencing as described by Christensen and colleagues, and performed at the DNA Sequencing Core Facility, University of Utah ^79^. In brief, 100 ng samples were prepared using an Illumina TruSeq DNA PCR-Free Library Prep, fragmented to a mean size of 200-250 bp, and sequenced on a NovaSeq Reagent Kit v1.5_150x150 flow cell, with a target read depth of 5M reads per sample. Due to the relatively short target DNA length (247 bp) and the viral DNA primer binding 84 bp upstream of the LTR (∼720 bp into the HIV-1 genome) generating a maximum integration junction less than 1Kb, sequencing the integration junctions presented inherent technical limitations. This sequencing protocol was optimized for samples larger than 1Kb, thus there were technical challenges for generating the expected thousands to hundreds of thousands of reads with our samples. As a result, we selected the samples that contained at the most contiguous integration junction reads for analysis. Therefore, two samples per NPS-DNA, unmodified-NCP, and H3K36me3-NCP, that met these criteria were analyzed. Next, fastq files were then filtered using SEAL to find reads that contained a 20 base kmer that mapped to the end of the 5’ LTR with at least 17/20 matches. These reads passing this filter were then mapped onto the nucleosome target sequences to generate sam and bam files. The Integrated Genomics Viewer (IGV) https://software.broadinstitute.org/software/igv/ was used to visualize the integration junctions ^143^. The integration junctions were tabulated by quantifying the continuous junction between the target DNA (in grey) and the viral DNA sequence (red, green, yellow, or blue), then plotted in excel.

## Supplemental Figures

**Supplemental Figure 1.**
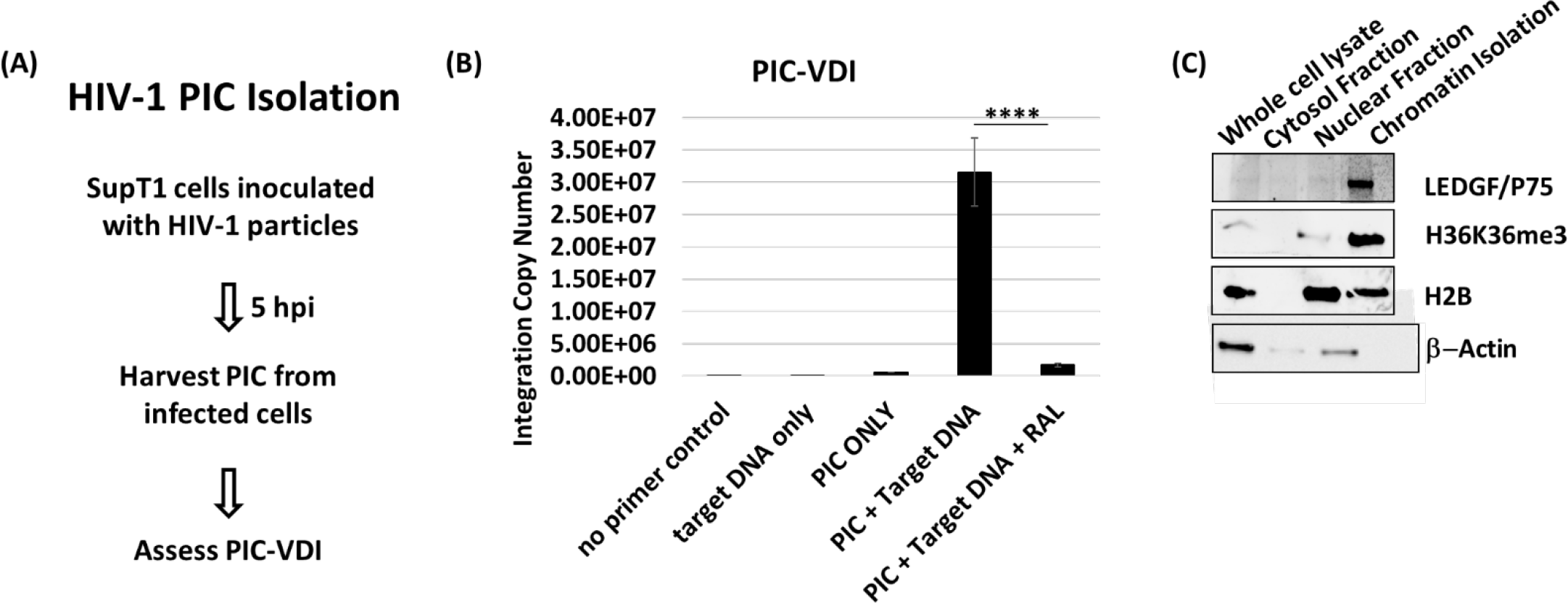
Virus infectivity and HIV-1 PIC isolation. (A) The isolation of HIV-1 PICs is shown as a diagram. First, HIV-1 virions were produced by transfecting HEK293T cells with the pNL4.3 HIV-1 molecular clone, collecting the supernatant 36-48 hours later, filtering the supernatant and treating with DNase I. Then HIV-1 pre-integration complexes (PIC) were isolated from SupT1 cells acutely infected with high tiger HIV-1 virions. The resultant PICs are assessed for viral DNA integration (VDI) using a nest-PCR assay. (B) HIV-1 PIC-VDI is measured by a nested PCR strategy that amplifies the DNA junction formed between the viral DNA and targeted substrate by the stand transfer activity of the HIV-1 integrase. The PIC-VDI is specifically inhibited by the integrase strand-transfer inhibitor raltegravir (RAL). The error bars were determined by the SEM and **** represents P < 0.0001. (C) The chromatin sample that was used as a substrate for PIC-VDI was probed by western blot analysis for the nuclear proteins associated with HIV-1 integration and cytoplasmic proteins.

**Supplemental Figure 2.**
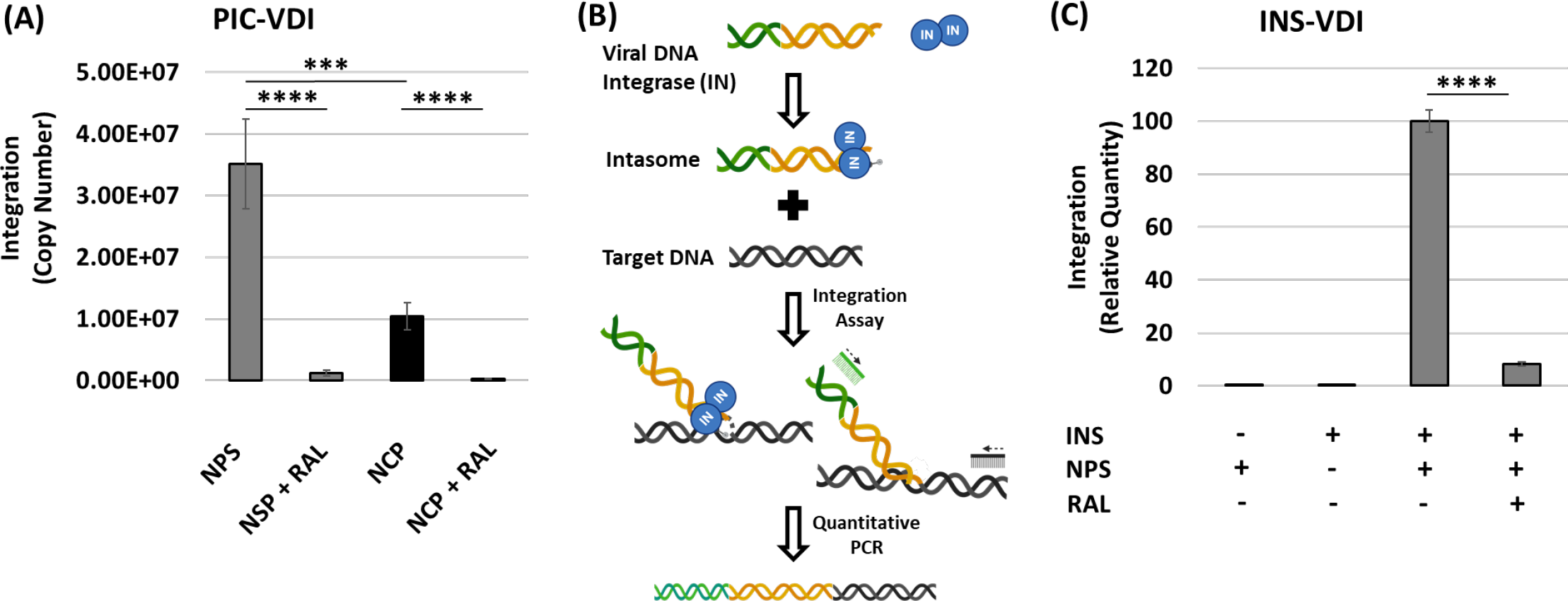
The recombinant NCP is a barrier to PIC-VDI and INS-VDI. (A) PIC- VDI was measured with the nucleosome positioning sequence (NPS) substrate and the nucleosome core particle (NCP) substrate, including RAL controls. The data is shown as the mean copy numbers of at least three independent PIC-VDI experiments. (B) A schematic of the intasome (INS) assembly. The INS was pre-assembled with the purified integrase protein and viral DNA sequences from the HIV-1 long-terminal repeat (LTR). Once assembled, the INS can carry out a DNA strand transfer reaction with any given target DNA. The INS-VDI is measured by quantitative PCR using primers designed for the INS donor DNA and the target DNA (acceptor DNA). (C) The INS-VDI was measured by qPCR with the NPS alongside the INS alone, substrate alone, or in the presence of the integrase inhibitor RAL. The error bars were determined by the SEM, and the *** represents P = 0.001 to 0.0001, **** P < 0.0001.

**Supplemental Figure 3.**
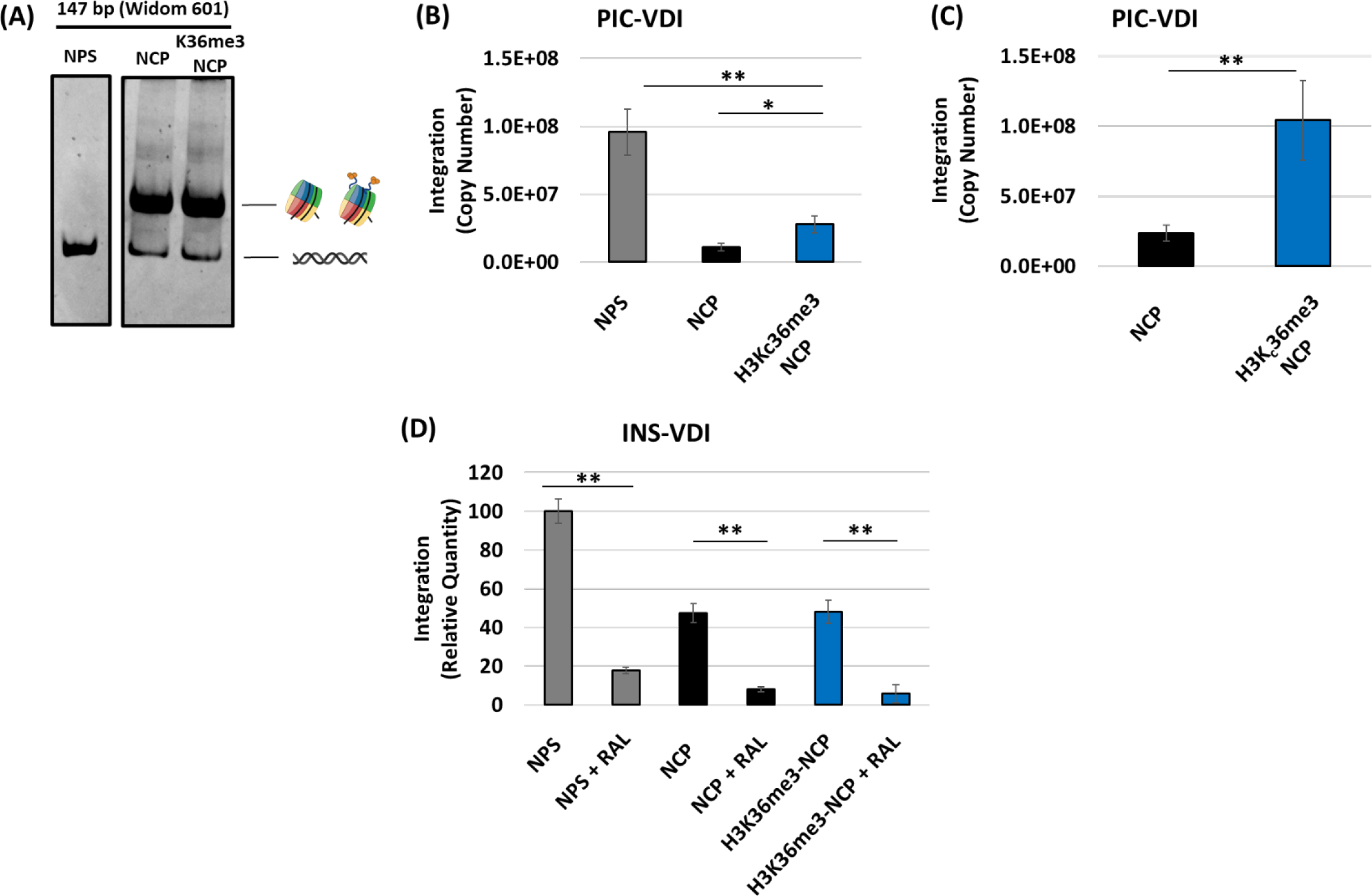
Comparative analysis of the PIC-VDI and INS-VDI with the NPS, NCP and H3K36me3-NCP. (A) The biochemically assembled NPS, NCP and H3K36Cme3-NCP were assessed by a 5% polyacrylamide gel electrophoresis (PAGE) and visualized with ethidium bromide staining. (B) Comparative analysis of the PIC-VDI with the NPS, NCP and H3K36me3- NCP was measured by nested qPCR, and the data is represented by the mean of viral DNA copy numbers from at least three independent experiments. (C) A direct comparison of the PIC-VDI with the NCP and H3K36me3-NCP is shown as the mean of copy numbers from independent experiments. (D) The INS-VDI was measured with the NPS, NCP and H3K36me3-NCP, including RAL controls. The error bars represent the SEM. * Represents P > 0.05 and ** P = 0.01 to 0.05.

**Supplemental Figure 4.**
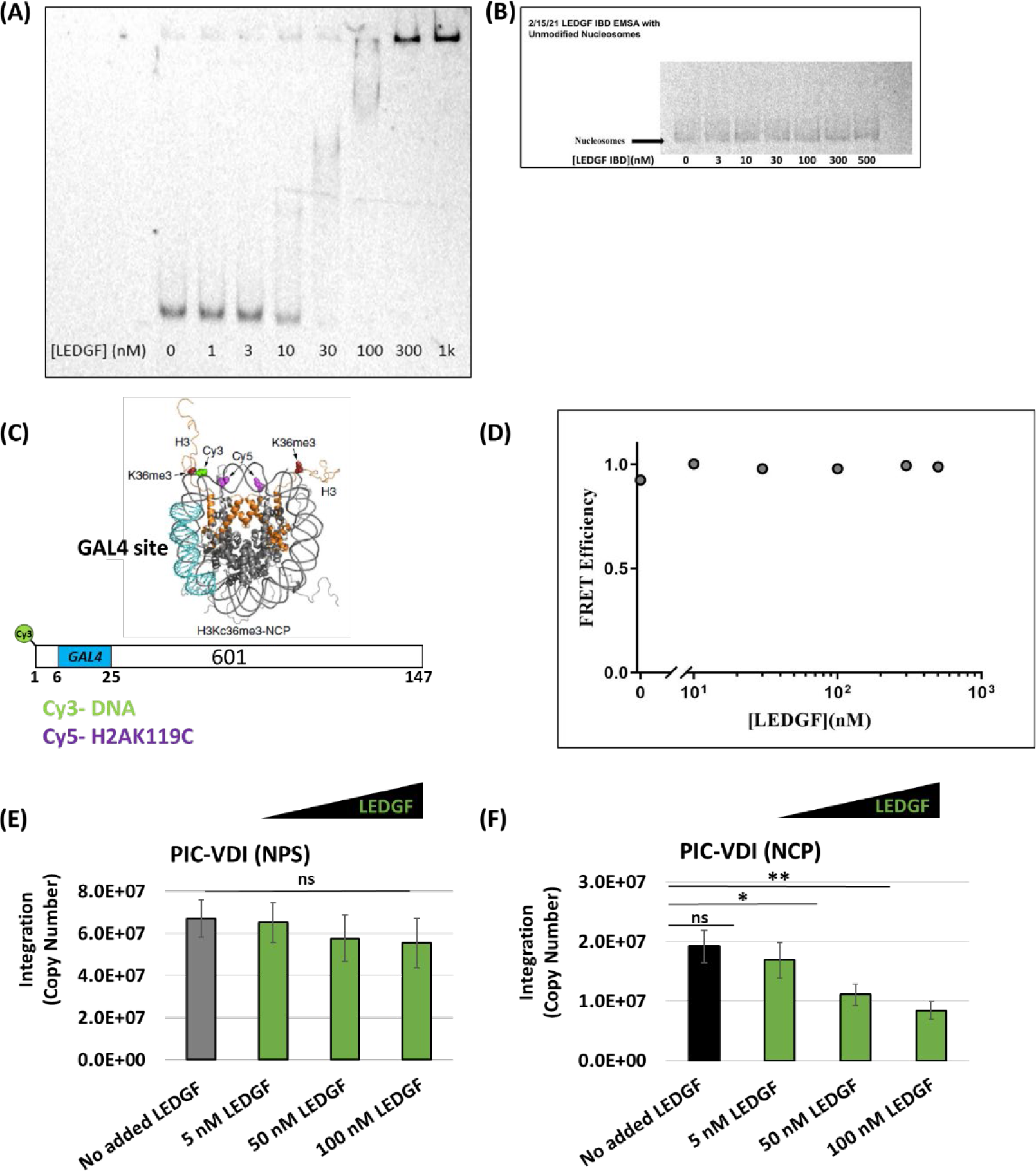
LEDGF/p75 binds the NCP and specifically reduces PIC-VDI with the NCP. (A) A schematic of the modified NCP for the FRET based binding assay is depicted. The ensemble FRET binding assay used NCPs labeled with Cy3 at the end of the NPS and a Cy5 at the H2AK119C position. A binding site for the transcription factor GAL4 was cloned into the NPS near the H3 tail protrusion site thru the nucleosomal DNA (6-25 bp) was designed to specifically determine the effect of LEDGF/p75 binding to the NCP. (B) The interaction between the NCP and LEDGF/p75 was assessed by EMSA. (C) Binding of the truncated mutant of LEDGF/p75, amino acids 341–429, that only contains the integrase binding domain (IBD) to the NCP was assessed for by EMSA. This truncated LEDGF/p75 lacks the AT-hook motifs nor the PWWP domain. (D) The FRET efficiency was measured in the presence of increasing amounts of LEDGF/p75. (E-F) The PIC-VDI with the NPS and the NCP were measured in the presence of LEDGF/p75 addition. These data are represented as the mean copy numbers of at least three independent experiments and the error bars represent the SEM. * Represents P > 0.05, ** P = 0.01 to 0.05, *** P = 0.01 to 0.001, P = 0.001 to 0.0001, and **** P < 0.0001.

**Supplemental Figure 5.**
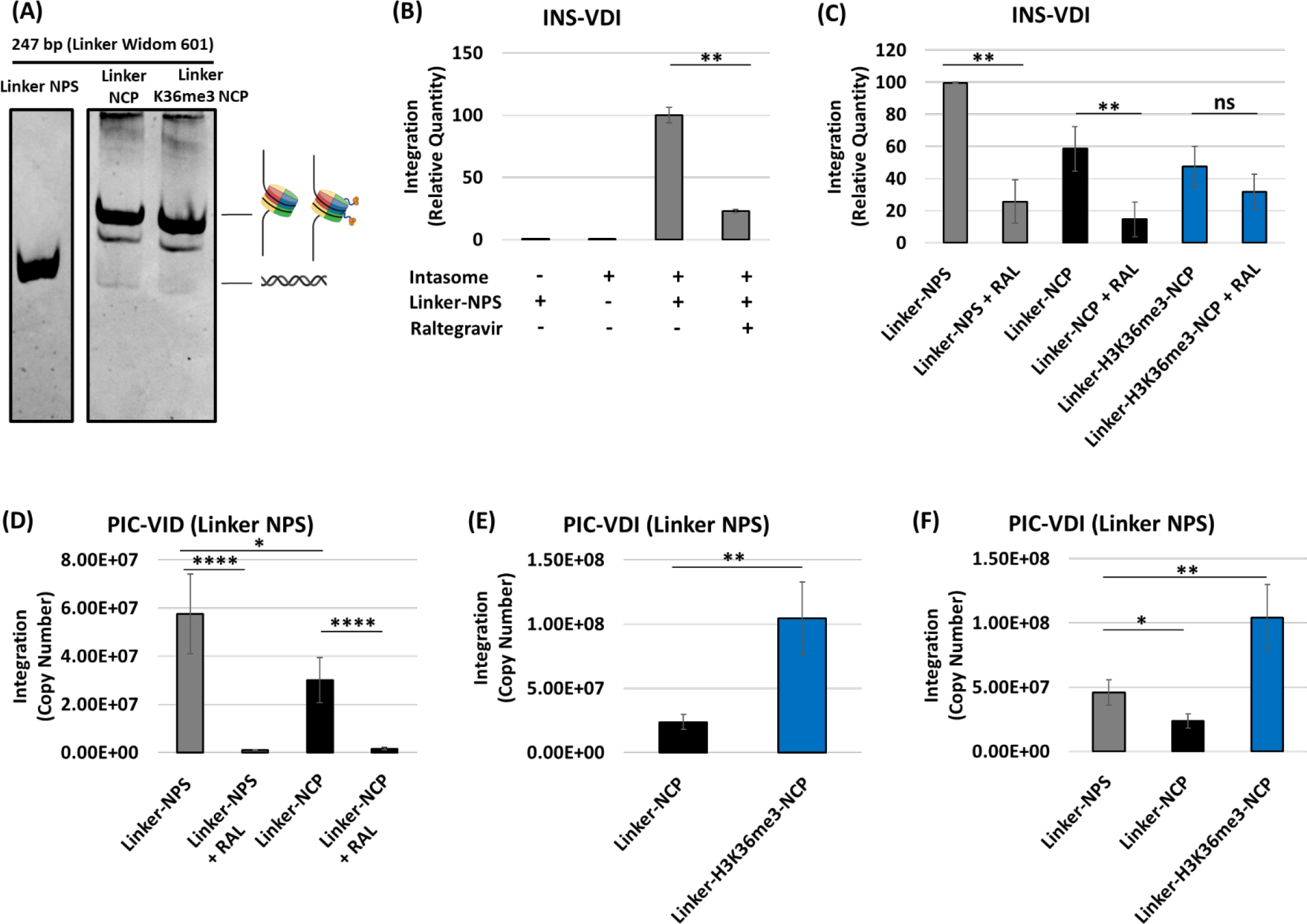
The linker-H3K36me3-NCP is an optimal substrate for PIC-VDI, despite the linker-NCP remaining a barrier to HIV-1 VDI. (A) The linker-NPS, linker-NCP, and linker-H3KC36me3-NCP were assessed by PAGE. (B) The INS-VDI was measured with the linker NPS, compared to the intasome only, target DNA only and RAL integrase inhibitor control. (C) The NPS, NCP and H3K36me3-NCP linker substrates were tested for INS-VDI along with RAL control assays. (D) The PIC-VDI was measured with the linker-NPS and linker-NCP substrate alongside RAL control assays. (E) The PIC-VDI was measured comparing the linker-NPS, to the linker-NCP and the linker-H3K36me3-NCP. (F) The linker-NPS, linker-NCP and linker- H3K36me3-NCP were compared as substrates for PIC-VDI. These results are shown as the mean copy numbers of at least three independent VDI experiments and the error bars represent the SEM. * Represents P > 0.05, ** P = 0.01 to 0.05, *** P = 0.01 to 0.001, P = 0.001 to 0.0001, and **** P < 0.0001.

**Supplemental Figure 6.**
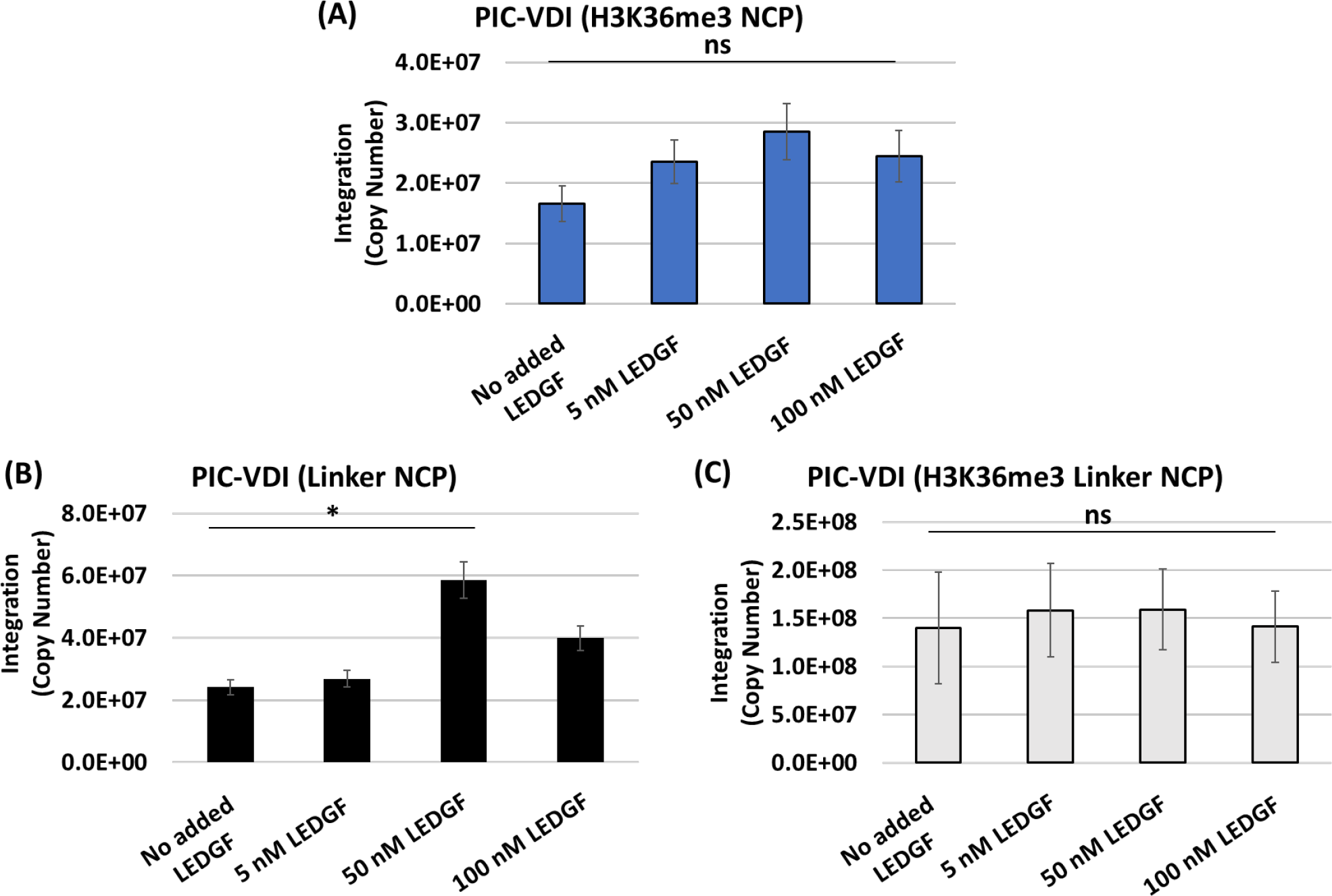
The LEDGF/p75 mediated inhibition of PIC-VDI is relieved by the H3K36me3 or the presence of linker DNA. PIC-mediated integration was measured with H3K36me3 NCPs in the presence of LEDGF/p75 addition. (A) PIC-VDI with H3K36me3-NCP was measured in the presence of LEDGF/p75 addition [5, 50 and 100 nM]. (B-C) PIC-VDI was measured with LEDGF/p75 addition to the linker-NCP and linker-H3K36me3-NCP substrates. These data are shown as the mean copy numbers of at least three independent experiments. The error bars were determined from the SEM and * Represents P > 0.05.

**Supplemental Figure 7.**
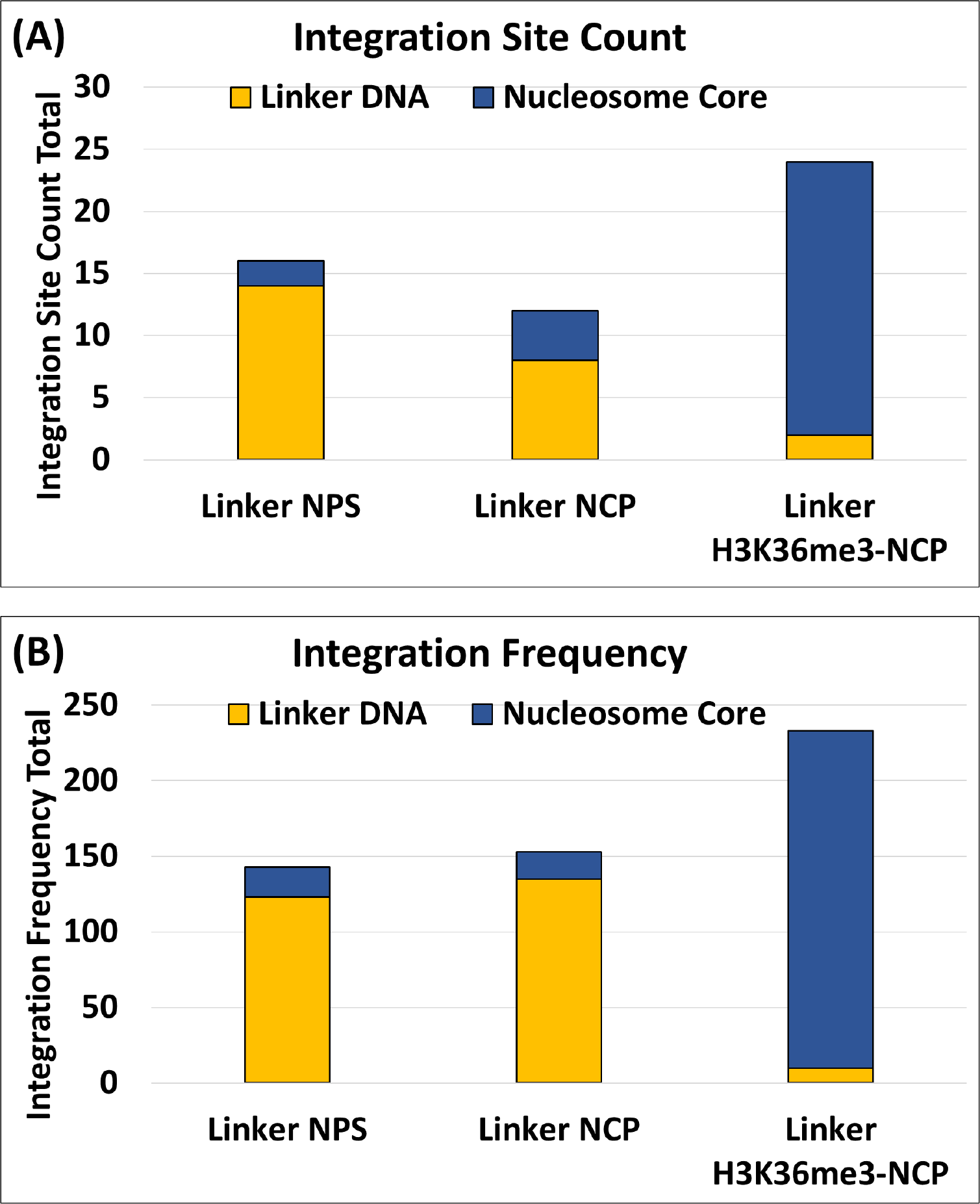
PIC-VDI within the linker-H3K36me3-NCP is overrepresented in the nucleosome core compared to the linker-NPS and linker-NCP substrates. To further study the HIV-1 integration preference within the linker DNA substrates, the DNA from the integration reactions were PCR amplified and subjected to next generation sequencing. Integration sites and frequency at those sites within the linker substrates was determined by quantifying the integration junctions. (A) Quantification of the integration sites within the linker substrates shows the number of integration events (sites) within the linker sequences (1-50 bp and 197-247 bp) and the nucleosome core sequence (50-197 bp). (B) The frequency of integration junctions at a particular site within the sequence was quantified as the integration frequency in the linker sequences and the nucleosome core sequence.

**Supplemental Figure 8.**
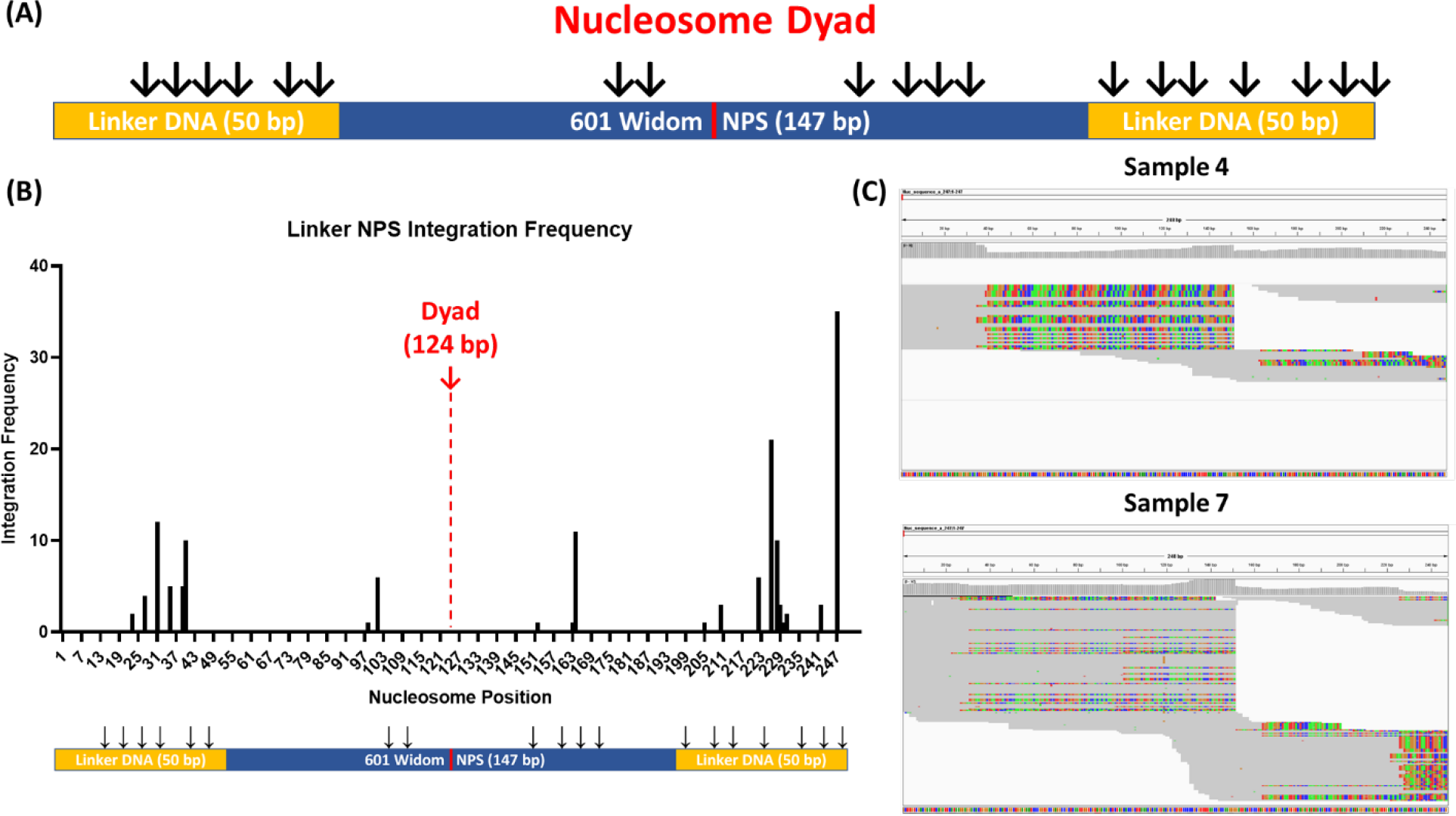
Distinct Integration sites with the linker NPS. The integration frequency within the linker-NPS was determined by quantifying the integration junctions along the substrate sequence. (A) A schematic representation of the integration sites within the linker NPS. (B) The integration frequency is plotted as a histogram for the linker NPS, showing the frequency of integration at particular sites of the substrate. (C) The raw IGV data showing the integration sites within the linker NPS substrate.

**Supplemental Figure 9.**
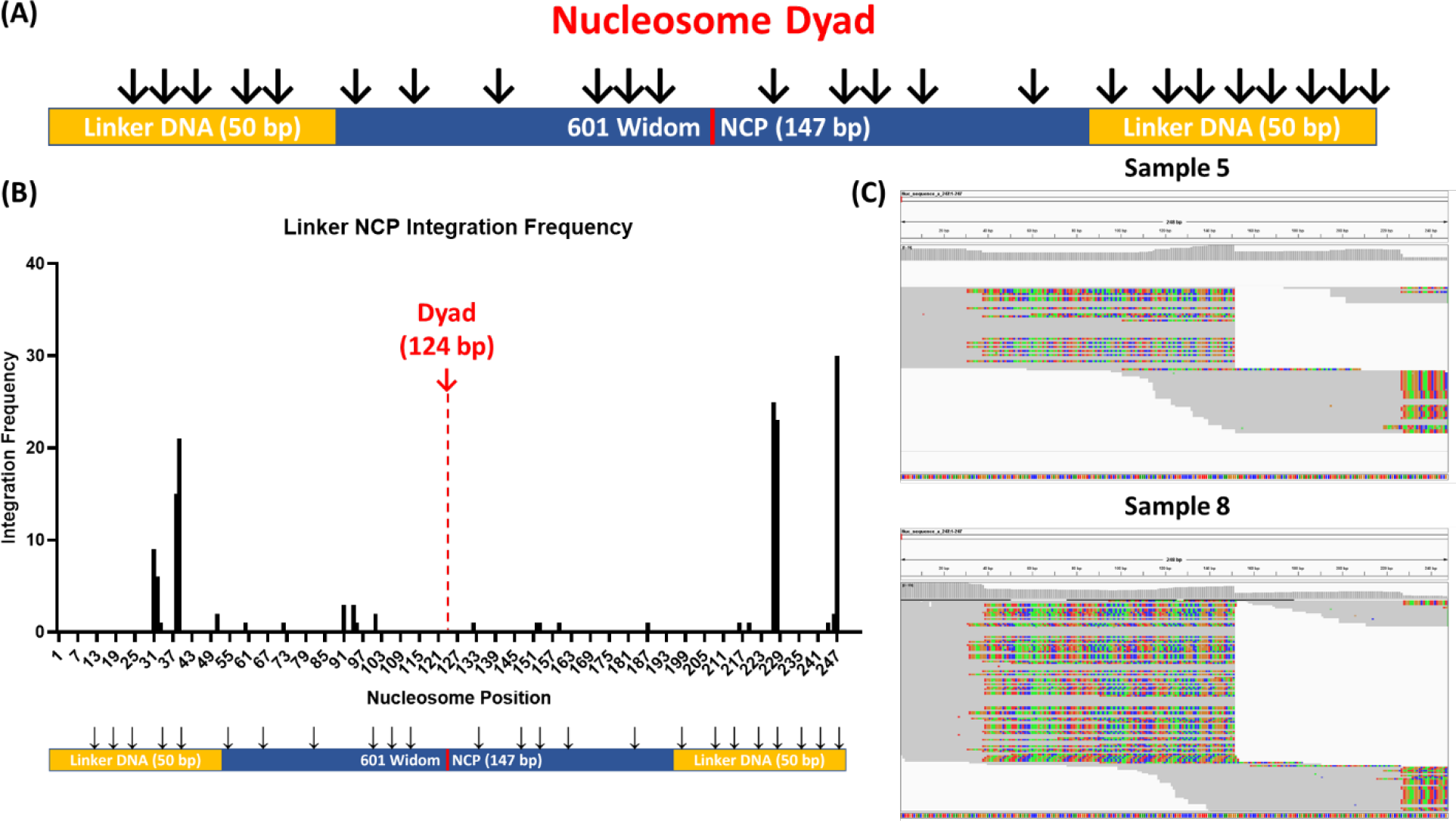
Distinct Integration sites with the linker NCP. The integration frequency with the linker-NCP was determined by quantifying the integration junctions along the substrate sequence. (A) A schematic representation of the integration sites within the linker NCP. (B) The integration frequency is plotted as a histogram for the linker NCP, showing the frequency of integration in specific sites of the substrate. (C) The raw IGV data showing the integration sites within the linker NCP substrate.

**Supplemental Figure 10.**
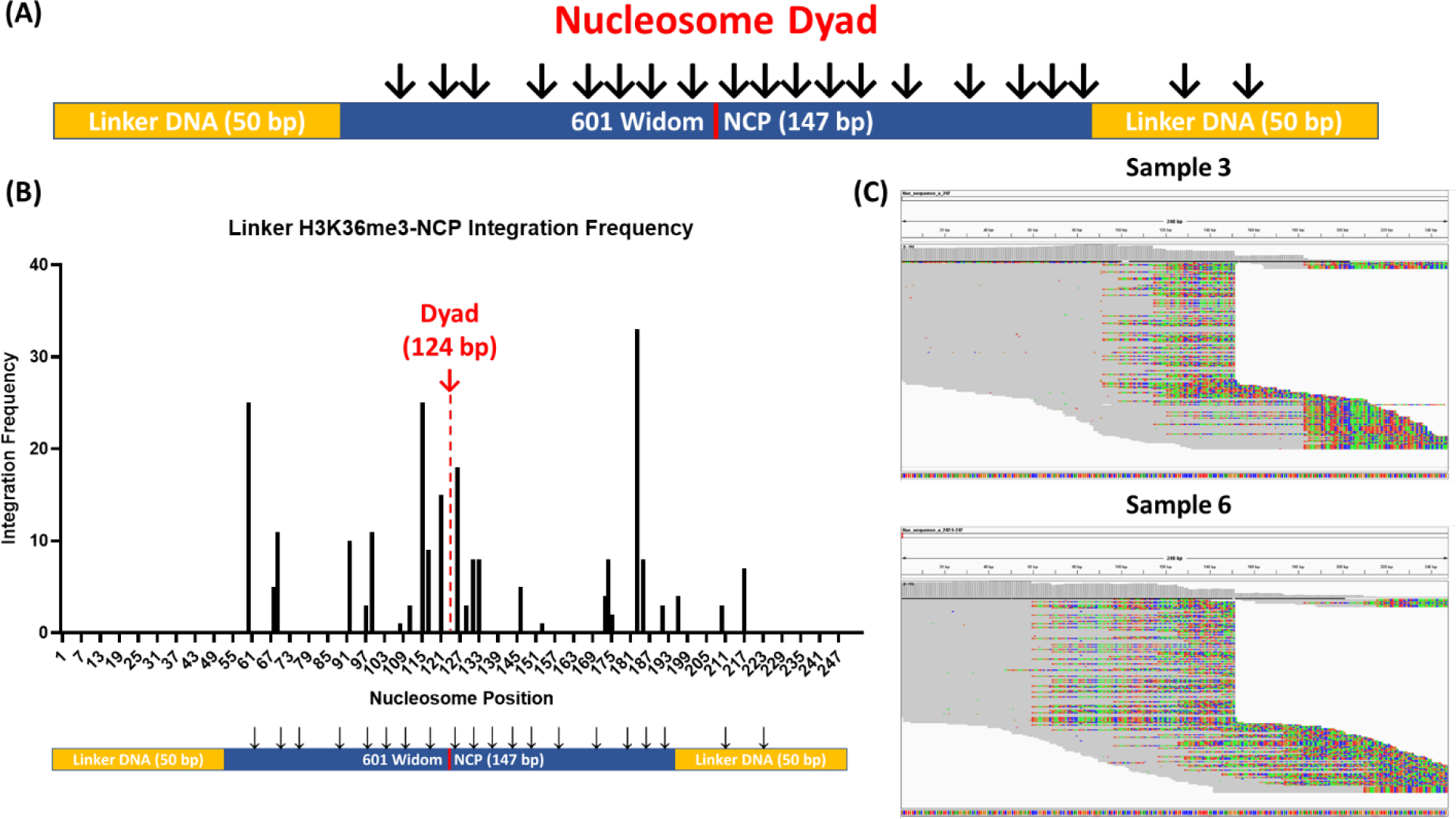
Distinct Integration sites with the linker H3K36me3-NCP. The integration frequency with the linker-H3K36me3-NCP was determined by quantifying the integration junctions along the substrate sequence. (A) A schematic representation of the integration sites within the linker H3K36me3-NCP. (B) The integration frequency is plotted as a histogram for the linker H3K36me3-NCP, showing the frequency of integration in specific sites of the substrate. (C) The raw IGV data showing the integration sites within the linker H3K36me3- NCP substrate.

**Supplemental Figure 11.**
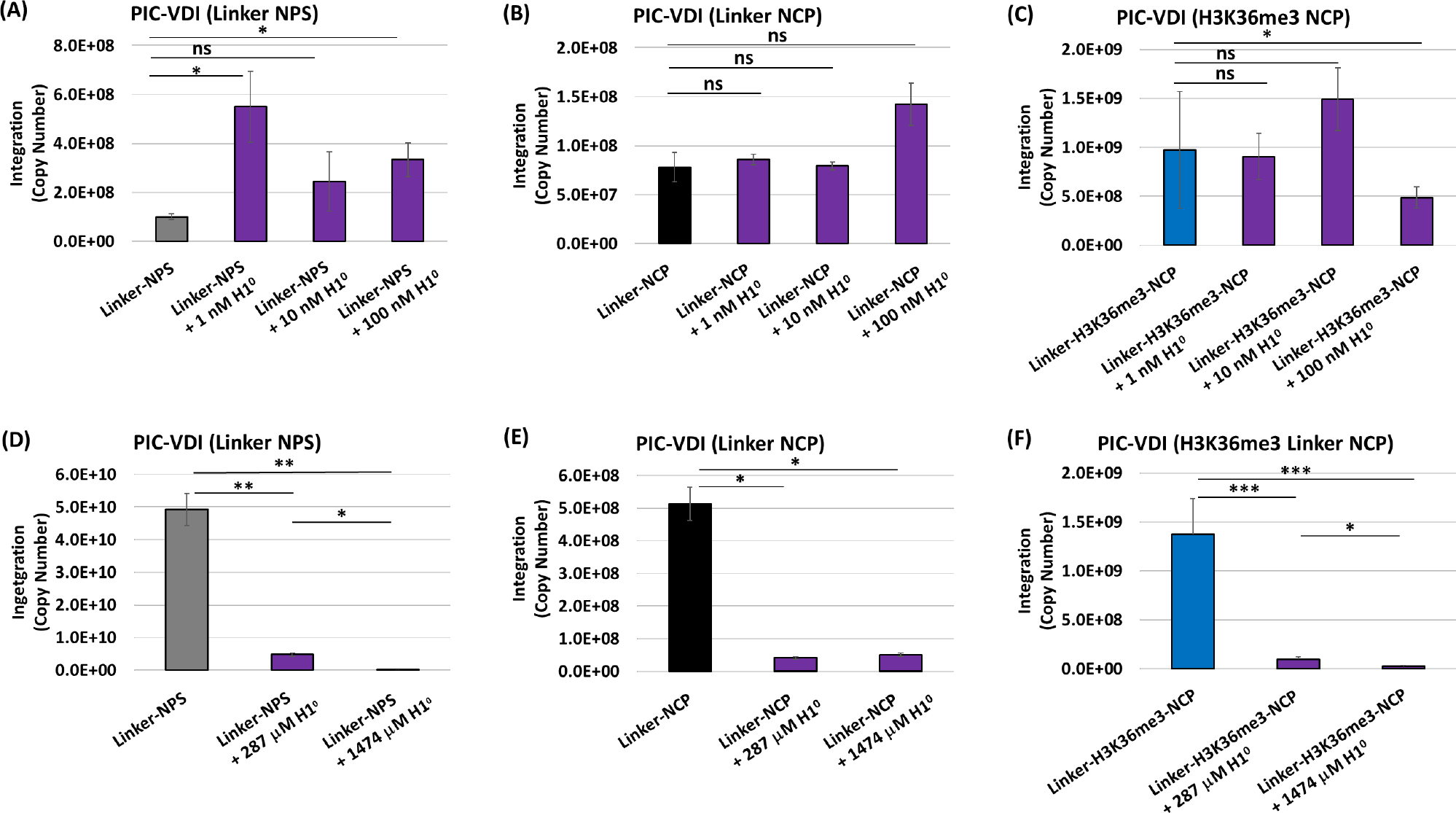
Linker histone H1 can inhibit PIC-VDI at saturating concentrations. The PIC-VDI with the linker substrates was measured with nM amounts of H1^0^ addition to the assay. (A-C) Non-saturating amounts of H1^0^ [1, 10 and 100 nM] were added to PIC-VDI with linker-NPS, linker-NCP, and linker-H3K36me3-NCP substrates. The data is represented as the mean of copy number from duplicate experiments. (D-F) The PIC-VDI with saturating amounts of H1^0^ [247 and 1474 µM] for the linker-NPS, the linker-NCP and the linker- H3K36me3-NCP are represented as the mean of viral DNA copy numbers from at least three independent experiments. For all data, the error bars indicate the standard error mean, and * represents P > 0.05, ** P = 0.01 to 0.05, *** P = 0.01 to 0.001, P = 0.001 to 0.0001, **** P < 0.0001.

**Supplemental Figure 12.**
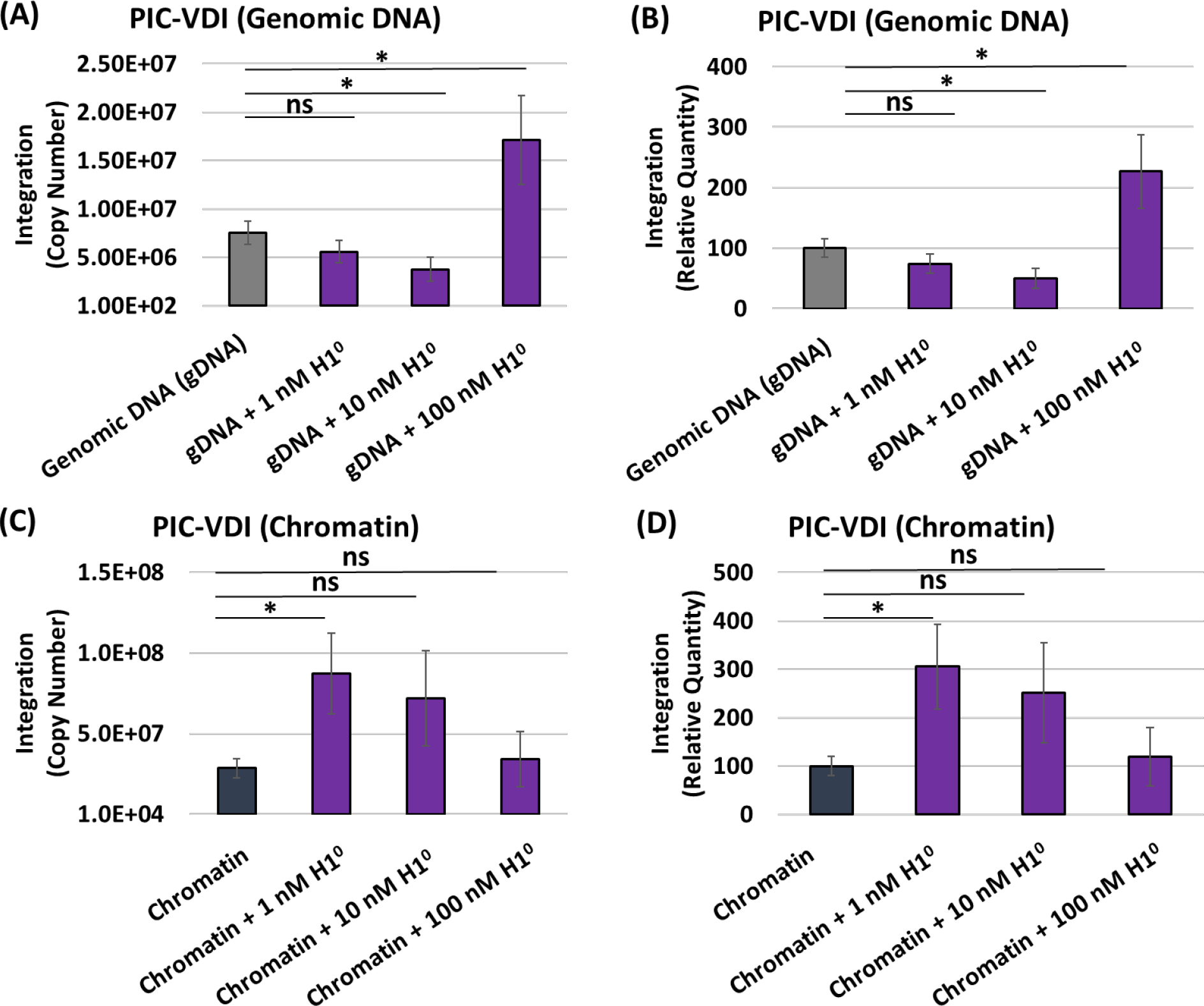
Non-saturating H1 addition to genomic DNA and chromatin differentially impacts PIC-VDI. The PIC-VDI with the genomic DNA (gDNA) and chromatin substrates was measured with nM amounts of H1^0^ addition to the assay. (A-B) The PIC-VDI with the genomic DNA in the presence of non-saturating amounts of H1^0^ is represented as the mean copy numbers of independent replicates and as the relative integration quantity. (C-D) The PIC- VDI with non-saturating amounts of H1^0^ incubated with chromatin is shown as the mean copy number and relative integration quantity. The error bars represent the standard error mean and * represents P > 0.05.

**Supplemental Figure 13.**
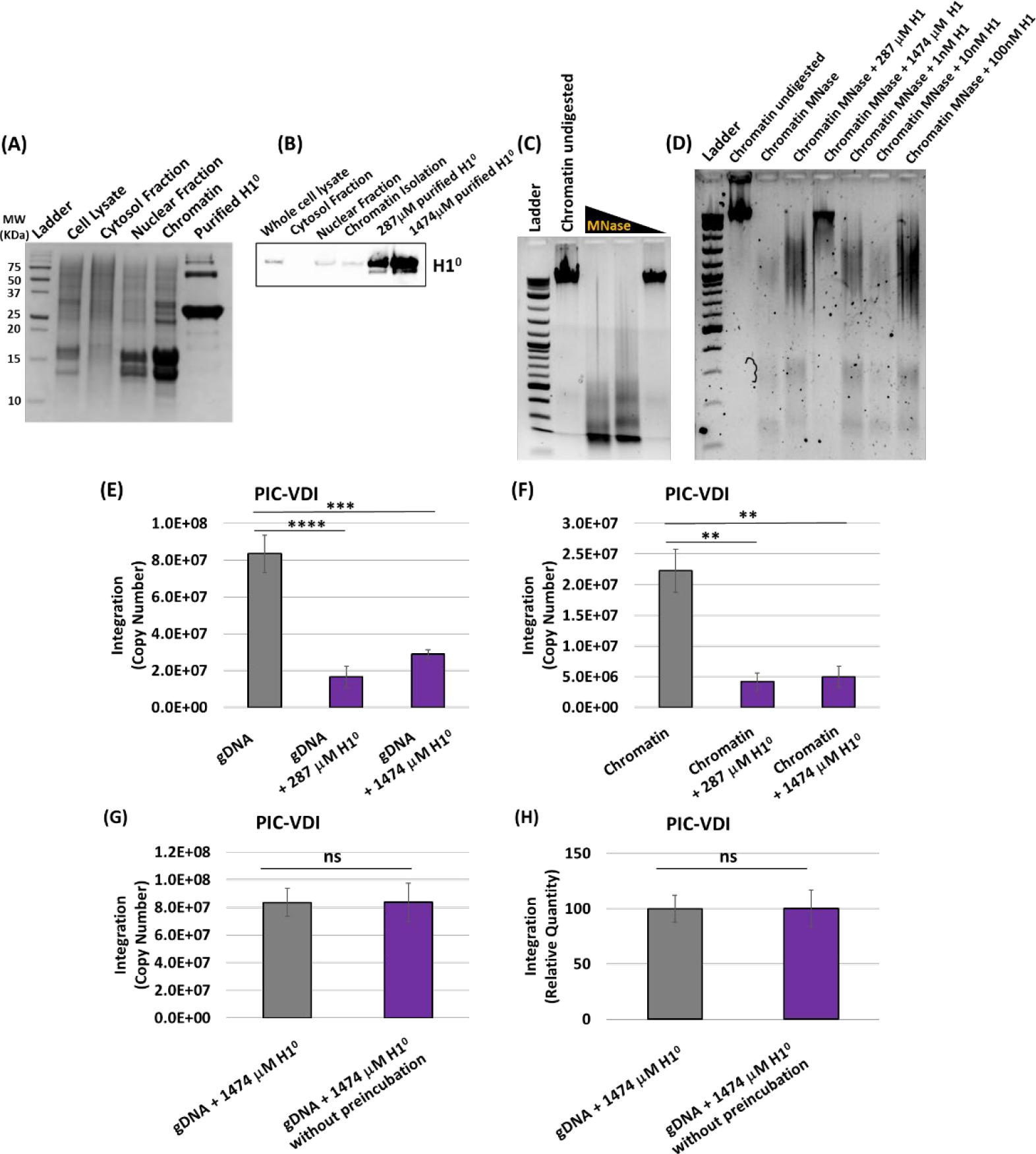
Saturation of H1 with chromatin and genomic DNA reduces PIC- VDI. (A) The purified recombinant H1^0^ (NEB) was assessed alongside the chromatin isolation fractions by Coomassie staining. (B) Western blot analysis of the chromatin isolation fractions and the saturating amounts of H1^0^ were probed for H1. (C) Chromatin samples isolated from HEK293T cells were assessed for nucleosome integrity by micrococcal nuclease (MNase) digestion at decreasing amounts. The resultant digested DNA was visualized by ethidium bromide staining of a 1.5 % regular agarose gel. (D) Chromatin samples pre-incubated with various amounts of H1^0^ were analyzed by MNase digestion and visualized by ethidium bromide staining of a 1.5% agarose gel. (E-F) The PIC-VDI was measured with genomic DNA and chromatin that were incubated on ice with 287 and 1474 µM of H1^0^ (saturating amounts). The µM concentrations reflect H1^0^ amounts that are 1:1 and 1:5 (w/w) of DNA substrate to H1^0^ protein. (G-H) A H1^0^ time-of-addition experiment with the saturating amount of H1^0^ is shown comparing the PIC-VDI with gDNA to an assay with H1^0^ added after the PIC was exposed to the substrate. The PIC-VDI data are represented as the mean of viral DNA copy numbers from at least three independent experiments and the error bars represent the standard error mean. The ** represents P = 0.01 to 0.05, *** P = 0.01 to 0.001, P = 0.001 to 0.0001, **** P < 0.0001.

**Table Supplementary 1.** Sequencing read counts for the PIC integration sites within linker DNA substrates.

| Sample Name | Sample # | Reads passing filter | Aligned reads |
| --- | --- | --- | --- |
| Linker<br>NPS | 4 | 268 | 57 |
|  | 7 | 440 | 194 |
| Linker<br>NCP | 5 | 188 | 80 |
|  | 8 | 620 | 250 |
| Linker<br>H3K36me3<br>NCP | 3 | 536 | 359 |
|  | 6 | 770 | 488 |

**Table Supplementary 2.**
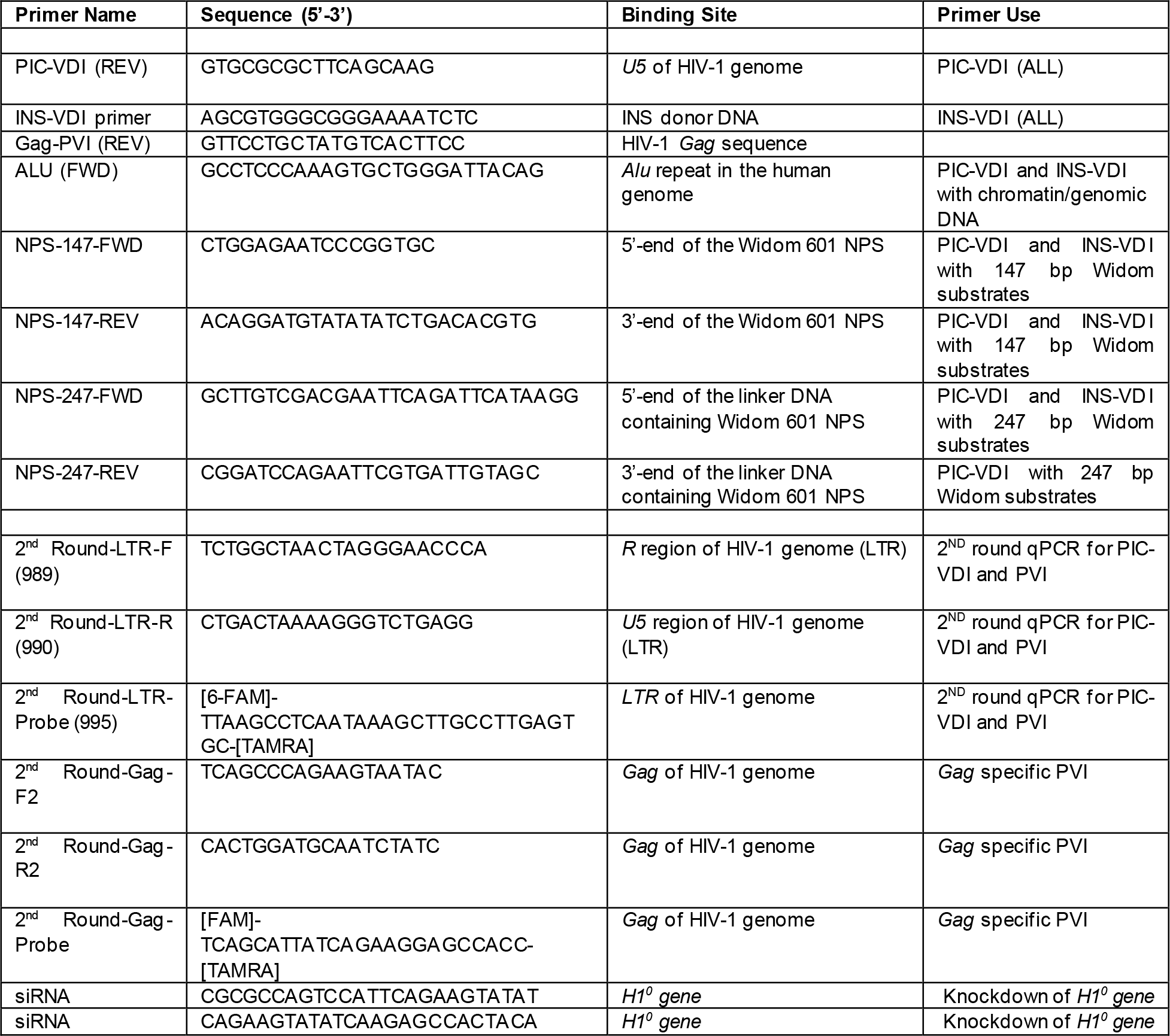
Primers used in this study.

## Notes

### Competing Interest Statement

The authors have declared no competing interest.

